# Molecular features underlying functional and structural interactions between diverse transcription regulatory elements in a chromosomal context

**DOI:** 10.64898/2026.02.16.706221

**Authors:** Yiyang Jin, Yuko Hasegawa, Zhou Zhou, Jessica Qin Lin, Alden King-Yung Leung, Haiyuan Yu, John T. Lis

**Affiliations:** Department of Molecular Biology and Genetics, Cornell University; Ithaca, NY, USA; Weill Institute for Cell and Molecular Biology, Cornell University; Ithaca, NY, USA; Department of Computational Biology, Cornell University; Ithaca, NY, USA

**Keywords:** Enhancer, Promoter, Transcription

## Abstract

Transcription is modulated by interactions among transcription regulatory elements (TREs), including promoters and enhancers, but their underlying interaction specificities remain opaque. Here, we develop a Chromatin-Integrated, landing-pad–based Enhancer Reporter Assay, CIERA-seq, that interrogates regulatory interactions between target promoters and distal TREs (dTREs) in a controlled genomic and chromatin context. We found both promoter-derived (pdTREs) and enhancer-derived-dTREs (edTREs) exhibit enhancer activities on target promoters with a specificity that depends on factors that dTREs recruit. pdTREs are generally enriched for factors involved in the core transcription process, whereas edTREs are enriched for pioneer factors, SWI/SNF chromatin remodelers, and histone acetyltransferase EP300. To further leverage the chromatin integration system, we applied deaminase-based footprinting to profile chromatin accessibility across individual fibers containing distinct target promoter–dTRE pairs. We found that different target promoters and dTREs exhibited distinct accessibility profiles, and that dTREs could modulate the accessibility of their cognate target promoters in a transcription factor and cofactor–dependent manner. Moreover, promoter accessibility was positively associated with transcriptional activity, linking dTRE-mediated changes in local chromatin accessibility to functional transcriptional output. Overall, our CIERA-seq assays and deamination-based footprinting assays support a model where dTREs activate target promoters constrained at different rate limiting steps in transcription by supplying complementary components of the transcription regulatory machinery.

## INTRODUCTION

Transcription is a dynamic and spatially organized process in which proximity between transcription regulatory elements (TREs), including promoters and distal TREs (dTREs) like enhancers, plays a central role in gene regulation^1,2^. A long-standing question in transcription regulation is whether promoters exhibit selective “compatibility” with specific enhancers. Early genetic studies established that enhancer–promoter interactions can exhibit substantial specificity, with individual enhancers preferentially activating certain promoters even when other promoters are located nearby^3,4^. Subsequent studies further demonstrated that core promoter elements, including the TATA box and downstream promoter element (DPE), can contribute to this specificity^5^. Plasmid-based assays across diverse systems, including Drosophila, mouse, and human cells, have produced mixed conclusions, with some studies reporting strong promoter specificity and others supporting broadly permissive enhancer-promoter regulation^6–11^. However, while plasmid-based assays are powerful tools enabling large-scale studies of regulatory interactions of enhancers and promoters, the chromatin structure of plasmids differs fundamentally from native chromatin. In the genome, DNA is packaged into nucleosomes, and chromatin accessibility is tightly regulated by intrinsic DNA features, histone modifications, and the activity of pioneer transcription factors (TFs) and chromatin remodelers^12–15^. Recent evidence indicates that, although a small fraction of transfected plasmids can assemble into relatively canonical chromatin-like structures, the majority either lack well-organized chromatin or exhibit aberrant nucleosome spacing^16^. This incomplete chromatin assembly can still allow detection of regulation at many steps in the transcription cycle but can compromise the study of regulatory steps that require alteration of chromatin. Consequently, regulatory principles governing promoter–dTRE interactions inferred from plasmid-based assays may not completely recapitulate how dTREs activate target promoters within an integrated chromosomal chromatin context^16,17^.

Another unresolved issue in transcriptional regulation arises from insights provided by high-resolution chromatin-conformation assays such as Hi-C and Micro-C, which have revealed multiple classes of TRE– TRE interactions, most prominently promoter–enhancer (P–E) and promoter–promoter (P–P) loops^18,19^. P– E interactions have been extensively characterized and are widely recognized as key mechanisms for achieving precise spatial and temporal control of gene expression^20^. In contrast, the functional significance of P-P interactions remains poorly understood ^19,21^. It is unclear the degree to which P–P loops merely reflect shared engagement with enhancer hubs or whether they play an active regulatory role by enabling promoters to directly influence one another’s transcription output ^6,22,23^. Some previous CRISPRi work has shown promoters can play potential regulatory roles like distal enhancers and regulate expression of other promoters^24^.

To overcome the limitations of plasmid-based assays and to investigate both promoter–enhancer compatibility and the regulatory significance of promoter–promoter interactions, we developed CIERA-seq, a Chromatin-Integrated, massively parallel Enhancer Reporter Assay that here enabled systematic interrogation of regulatory interactions between 16 target promoters and approximately 350 dTREs spanning diverse regulatory features^25,26^. Target promoters differ markedly in their intrinsic chromatin accessibility and basal transcription activity, and these intrinsic properties, likely a result of the differential enrichment of transcription machinery (TFs and co-activators) at these promoters, strongly shape their responsiveness to dTREs. Notably, we observed that many target promoters can be activated over a large dynamic range not only by enhancer-derived-dTREs (edTREs) but also by other promoter-derived-dTREs (pdTREs). This widespread promoter–pdTRE activation suggests that promoter-promoter interactions play a more prominent and direct regulatory role at endogenous loci than previously appreciated. We also identified that pdTREs and edTREs are enriched for different categories of transcription machinery, which may explain their difference in regulatory activity. We further leveraged the chromatin-integrated nature of CIERA-seq by applying deaminase-based footprinting at the landing-pad locus, enabling simultaneous measurement of chromatin accessibility across paired promoter–dTRE constructs. We then combined this framework with targeted mutational analyses to dissect the contributions of specific transcription factors to dTRE-mediated promoter regulation. Integrating the CIERA-seq and footprinting datasets, we propose a mechanistic model in which promoters that are deficient in specific components of the transcriptional machinery are preferentially activated by dTREs capable of supplying or recruiting those missing components. These findings support a model in which sharing transcriptional machinery between dTREs and promoters is a key determinant of the specificity of dTRE-mediated promoter activation^13^.

## RESULTS

### Design of CIERA-seq

To investigate how target promoters respond to activation by diverse dTREs in a native chromatin context, we developed CIERA-seq. Using this landing-pad platform, we quantified the activities of more than 350 dTRE candidates paired with 16 target promoters in K562 cells, yielding around 5,000 target promoter– dTRE combinations. We first generated a K562 cell line harboring a Bxb1 recombinase landing pad with constitutive BFP expression inserted at the eNMU safe-harbor locus^26^. In our recent work, we demonstrated that, in the landing-pad cell line in which the endogenous eNMU elements are deleted, the entire TAD encompassing the eNMU locus lacks detectable chromatin accessibility and transcriptional activity from neighboring dTREs. This neutral regulatory environment therefore provides an ideal genomic context for performing MPRA-based assays by minimizing potential confounding effects from surrounding cis-regulatory elements. Site-specific integration of a target promoter–EGFP–dTRE construct replaces the BFP cassette, such that recombined cells lose BFP and express EGFP under the control of the integrated promoter–dTRE pair (where dTRE refers to distally located element downstream of EGFP, be it enhancer-derived or promoter-derived) (Figure 1A). For a given target promoter, the expectation is that dTREs with stronger regulatory activity drive higher EGFP expression than weaker ones (Figure 1A). Cells containing different target promoter–dTRE combinations were collected into seven EGFP expression bins, and subjected to sequencing of both the target promoter barcodes (PIDs), identifying the target promoters, and the dTRE sequences from each bin^27^. This approach enabled us to derive bin-specific expression distributions for every target promoter–dTRE pair and to calculate an activation score based on weighted EGFP bin values (Figures 1B and S1A). Scores were reproducible across three biological replicates (Figure S1B-C).

**Figure 1.**
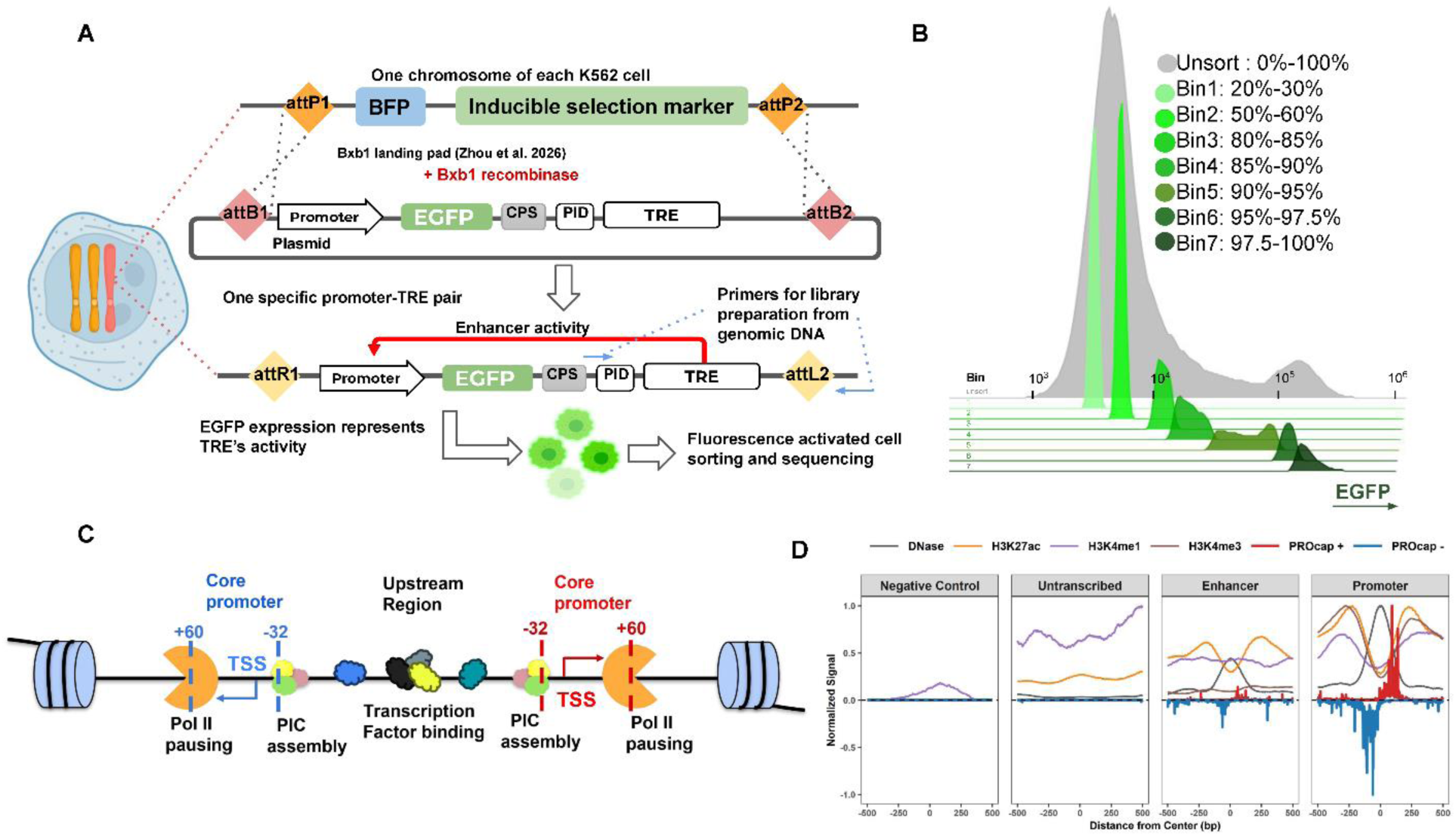
Chromatin-integrated assay to quantify promoter–dTRE regulatory interactions. (A) Schematic of the chromatin-based landing pad assay used to measure the regulatory activity of >350 transcriptional regulatory elements (dTREs paired with 16 promoters, CPS refers to cleavage and polyadenylation sites and PID refers to promoter identity barcode). (1) A Bxb1 landing pad is inserted at the enhancer locus of the NMU gene (chr4:55,729,891-55,730,846) on one chromosome to generate a K562 landing pad cell line. (2) dTREs are cloned into an EGFP reporter vector containing Bxb1 recombination sites together with distinct promoters and corresponding promoter barcodes (PIDs). (3) Vectors are co-transfected into K562 cells with a Bxb1 recombinase expression plasmid. (4) Cells are cultured for 1 week and sorted for loss of BFP signal, which indicates successful integration. (5) Cells are cultured for >3 weeks and sorted into bins based on EGFP expression. (6) Genomic DNA f rom each bin is extracted and used to generate Illumina sequencing libraries containing PID and dTRE sequences with primers shown (blue arrows) in the illustration. (7) Sequencing data are analyzed as described in Figure S1A. (B) Representative FACS profile of K562 cells. Each cell has a single promoter–dTRE pair integrant and gets sorted into seven bins based on EGFP expression. Gray indicates the unsorted population; colored peaks represent individual bins. Bin boundaries are defined by percentiles of the EGFP expression distribution across the population. (C) Definition of pdTREs and edTREs used in this study. Both pdTREs and edTREs are defined by divergent PRO-cap signals and contain two divergent core promoters, each defined by where the Pol II PIC begins at-32 bp and extends through the pause region to +60 bp f rom the TSS. The region between the two core promoters serves as an upstream regulatory region enriched for transc ription factor binding. pdTREs are defined as dTREs overlapping annotated transcription start sites of protein-coding or lncRNA genes, whereas edTREs are located in intergenic regions or within introns. (D) Profiles of PRO-cap, DNase-seq, and ChIP–seq signals for H3K27ac, H3K4me1, and H3K4me3 across dTREs of different element types.

We defined dTREs, both edTREs and pdTREs, using divergent PRO-cap transcription start sites and extended each element by ±60 bp to capture features required for transcription initiation, pausing, and release (Figure 1C)^11,28–31^. Most dTREs included in this analysis were previously identified using ChromHMM annotations and divergent transcription as complementary markers of regulatory activity and were subsequently characterized across multiple plasmid-based MPRA platforms^6,11^. These prior measurements revealed a broad range of regulatory activities, from weakly active to highly active, depending on the promoter with which each dTRE was paired (Figure S2A). This selection strategy therefore served two purposes: it enabled direct comparison of dTRE activity between plasmid-based and chromatin-integrated contexts, and it increased the likelihood of capturing a broad dynamic range of regulatory activities following genomic integration. To facilitate a balanced comparison between pdTRE and edTRE classes, we also included approximately equal numbers of pdTREs and edTREs in the analysis (Figure S2B). Although pdTREs and edTREs share similar regulatory architectures, pdTREs are distinguished by their gene-proximal context and the frequent presence of 5′ splice sites that stabilize pre-mRNA, whereas edTREs are typically distal or intronic and generate unstable enhancer RNAs (Figure 1C)^32,33^. pdTREs and edTREs differ in their chromatin states and transcription levels. pdTREs exhibited higher PRO-cap, DNase-seq, H3K27ac, and H3K4me3 signals than edTREs, as seen previously^32^, whereas H3K4me1, a frequently used enhancer mark, did not robustly distinguish between the two classes (Figure 1D)^32^. In addition to open reading frame (ORF)-derived negative controls previously shown to exhibit no detectable enhancer activity in plasmid-based reporter assays, we also included a set of “untranscribed” dTREs that lacked detectable PRO-cap signal but were annotated as enhancers based on ChromHMM and ENCODE cCREs and enriched for H3K4me1 (Figures 1D and S2B-C)^14,34,35^.

### Diverse compatibilities of distal TRE-promoter activation

The 16 target promoters were selected to span a broad range of endogenous transcription states, as defined by PRO-seq at their native loci^28^ (Figures S2D and S2E). To examine how intrinsic promoter properties influence their responsiveness to dTREs, we included both highly expressed promoters at their native genomic locus, such as *MYC* and *GAPDH*, and promoters that are transcriptionally silent in K562 cells but active in other cell types, including *HNF1B*, *ADGRG5*, *BIN2*, and *MNDA* (hereafter referred to as untranscribed promoters)^36^. For each target promoter, the distribution and mean score obtained when paired with negative control elements provided a measure of its intrinsic activity at the integration locus (Figure 2A). Target promoters exhibited wide variation in intrinsic activity and showed only a modest positive correlation to PRO-seq transcription levels at their endogenous loci (Figure 2B). The relatively modest correlation between promoter activity at the endogenous locus and activity following integration into the neutral landing-pad locus is expected, as endogenous transcriptional output is determined not only by the intrinsic regulatory capacity of the promoter itself but also by inputs from the surrounding cis-regulatory environment. The marked reduction in *MYC* promoter activity at the eNMU safe harbor locus, despite high endogenous transcription, may reflect its separation from the numerous strong enhancers at its native locus (Figures 2A and S2D)^20^. Conversely, the high landing-pad activity of the *CNBP* promoter may reflect relief from native-locus silencing, consistent with findings in mouse^37^. These observations suggest that endogenous promoter activity reflects both promoter-intrinsic capacity and regulatory input from neighboring dTREs, whereas activity in the standardized landing-pad context provides a measure of intrinsic promoter capacity (Figures 2B and S2D–E).

**Figure 2.**
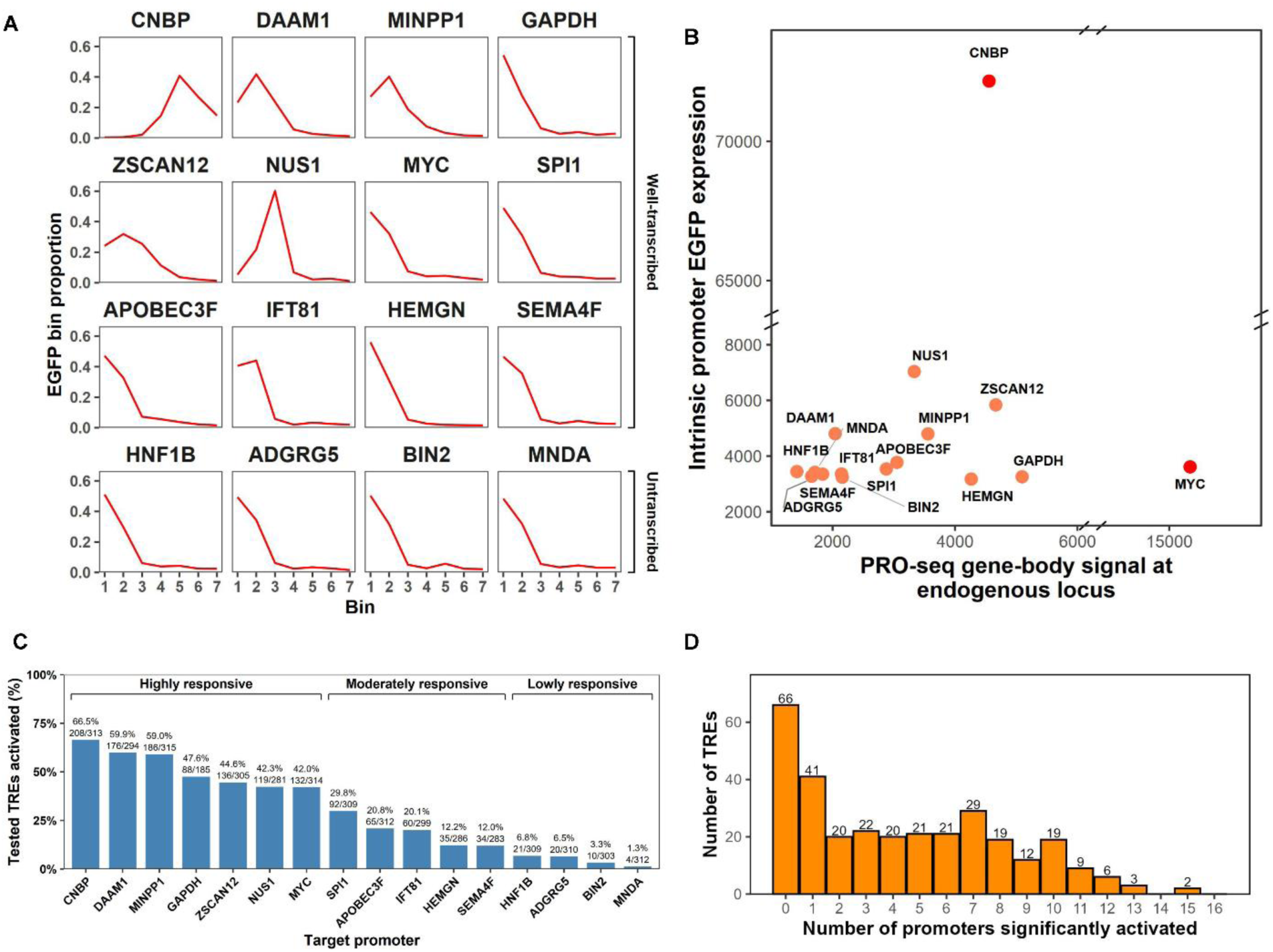
Different target promoters show different activity at the landing pad locus. (A) Mean normalized EGFP-bin proportions (red line) across seven expression bins for each of the 16 promoters paired with ORF negative controls, reflecting intrinsic promoter activity at the integration locus. Well transcribed and untranscribed are defined based on the activity of these target promoters at their native locus. (B) Correlation between intrinsic promoter activity at the integration locus calculated based on distribution across the EGFP expression bins and PRO-seq gene body signal per kb at the native genomic locus. (C) Percentage of successfully cloned dTREs (excluding ORF controls) that significantly activated each of the 16 target promoters. (D) Distribution of the number of target promoters significantly activated by individual dTREs. dTREs were included when at least 11 target promoters were testable.

For each target promoter, we counted how many dTREs can significantly increase the expression of EGFP from the target promoter compared to ORF-derived negative controls. As expected, some strong dTREs strongly shift the distribution of each promoter towards the right side (Figure S2F), consistent with the experimental design. In general, well-expressed promoters at their native loci are activated by a larger number of dTREs than untranscribed promoters. Promoters differed markedly in their responsiveness to dTRE input, revealing three broad classes: 1) well-expressed promoters that are highly responsive to dTREs, including *CNBP*, *DAAM1*, *MINPP1*, *GAPDH*, *ZSCAN12*, *NUS1*, and *MYC*; 2) well-expressed promoters that are moderately responsive to dTREs, including *SPI1*, *APOBEC3F*, *IFT81*, *HEMGN*, and *SEMA4F*; and 3) untranscribed promoters that are lowly responsive to dTREs, including *HNF1B*, *ADGRG5*, *BIN2*, and *MNDA* (Figure 2C). dTREs also varied widely in their specificity of enhancer activity, with some activating up to 15 of the 16 target promoters tested, whereas others activated only one or none, consistent with previous studies^8^ (Figures 2D and S2G).

### CIERA-seq and plasmid-based QUASARR-seq reveal distinct patterns of distal TRE activity across target promoters

We next compared promoter–dTRE pairs assayed in both the chromatin-integrated CIERA-seq system and the plasmid-based QUASARR-seq assay. Although the same promoter–dTRE combinations were tested in both contexts, the two assays revealed distinct distributions and dynamic ranges of regulatory activity (Figure S3A). For example, dTREs paired with the *GAPDH* promoter exhibited a substantially broader range of regulatory activities in CIERA-seq than in QUASARR-seq. In contrast, dTREs paired with the *MYC* promoter showed comparable activity ranges across the two assays. Conversely, dTREs paired with the *APOBEC3F* promoter exhibited a narrower range of activities in CIERA-seq relative to QUASARR-seq. Thus, the effects of individual dTREs on target promoters can differ between episomal and chromatin-integrated environments, and the magnitude and direction of these differences are themselves promoter dependent. These observations highlight the importance of chromatin context in a neutral safe harbor locus for evaluating the intrinsic behavior of promoter–dTRE regulatory interactions.

### Both enhancer-derived distal TREs and promoter-derived distal TREs show enhancer activity

We previously showed that both promoters and enhancers exhibit enhancer-like activity in a plasmid-based assay, but whether this activity is retained in chromatin remained unclear ^6^. Across target promoters, both pdTREs and edTREs exhibited substantially stronger enhancer activity than untranscribed elements and negative controls (Figure 3A), with pdTREs showing broader compatibility with target promoters than edTREs (Figure S2G). Target promoters varied markedly in their responses to dTRE input. Highly responsive promoters, including *CNBP*, *DAAM1*, *MINPP1*, *GAPDH*, *ZSCAN12*, *NUS1*, and *MYC*, were activated more strongly and by a broader range of pdTREs than edTREs. Moderately responsive promoters, including *SPI1*, *APOBEC3F*, *IFT81*, *HEMGN*, and *SEMA4F*, responded similarly to the two groups (Figure 3A). In contrast, lowly responsive untranscribed promoters, including *HNF1B*, *ADGRG5*, *BIN2*, and *MNDA*, were activated by only a limited subset of dTREs (Figure 3A). This pattern suggests that certain dTREs possess specific regulatory features that enable them to activate otherwise untranscribed target promoters more effectively than other dTREs.

**Figure 3.**
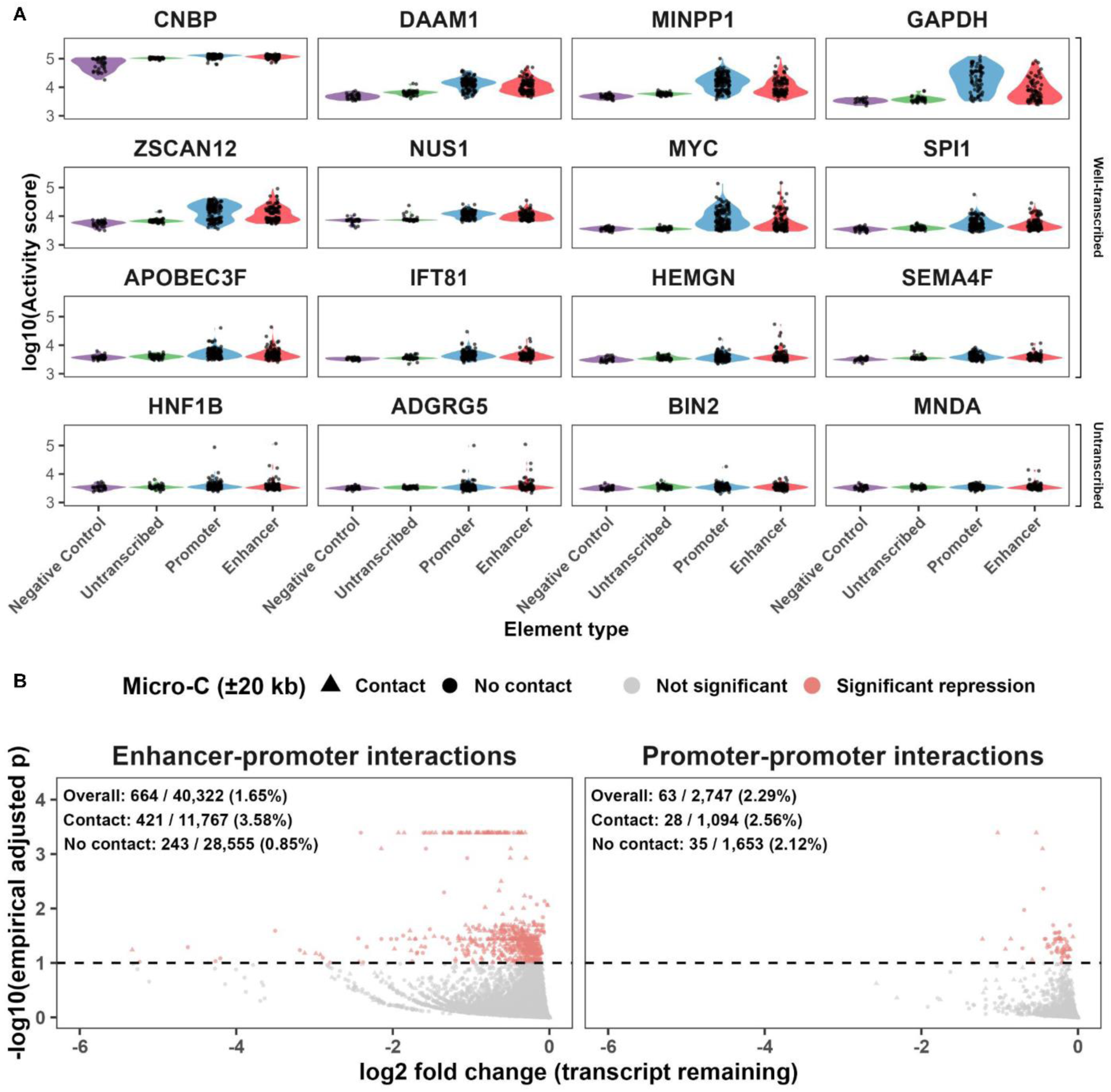
Different target promoters show different levels of activation by different classes of dTREs. A. Violin plot showing the distribution of EGFP expression activity scores for the 16 target promoters when paired with dTREs of different element types. B. Volcano plots showing changes in gene expression associated with guide RNAs groups targeting enhancer or promoter regions, noting whether they have or do not have micro-C contacts.

These results suggest that pdTREs can function as regulatory elements in the CIERA-seq setting. Given previous evidence that promoters can engage in promoter–promoter chromatin interactions and potentially regulate the expression of other genes, we sought to investigate whether promoters also exhibit regulatory activity in their native genomic contexts^6,22,23^. To address this question, we reanalyzed publicly available CRISPRi candidate enhancer perturbation data to assess the potential regulatory effects of promoter silencing on genes other than those directly associated with the targeted promoters (Figure 3B)^24^. To reduce potential indirect trans effects, we excluded gRNA groups targeting genes with known or putative roles in transcriptional regulation. We found that 63 of 2,747 promoter-targeting gRNA groups (2.29%) were associated with significant downregulation of other genes, a modestly higher proportion than that observed for enhancer-targeting groups (664 of 40,322; 1.65%). However, the magnitude of downregulation was generally smaller following promoter silencing than enhancer silencing, indicating that the frequency and strength of these regulatory effects differed between the two classes. To examine whether these effects were associated with physical interactions, we incorporated Micro-C data^38^. Both promoters and enhancers were more likely to exhibit regulatory effects when they contacted the corresponding target promoters, suggesting that chromatin interactions help shape endogenous dTRE activity. Together, these findings support a role for promoters in regulating neighboring genes, consistent with their enhancer activity in CIERA-seq, although residual trans effects cannot be excluded.

### Enhancer-derived distal TREs and promoter-derived distal TREs are enriched for different TFs and coactivators

In our model, most dTREs are comprised of a central region containing transcription factor binding sites flanked by a pair of divergent core promoters that support RNA polymerase II (Pol II) binding, initiation, pausing, and pause release^11,32^ (Figure 1C). In the CIERA-seq system, the dTRE is positioned in relatively close proximity (∼ 1.4 kb) but downstream of the target promoter, leading us to hypothesize that the dTREs activate the target promoter by sharing transcription machinery with it ^39,40^ (Figure 1A). To examine how dTRE-associated transcription machinery relates to dTRE’s enhancer activity, we analyzed ChIP-seq profiles at the endogenous loci of the tested dTREs and used 153 selected transcription factors and cofactors to characterize differences among dTRE classes ^35,41,42^. Clustering based on TF occupancy partitioned the 320 non-ORF TREs into seven final groups defined by characteristic transcription factor occupancy patterns (Figure 4A, selected TFs with known activating roles; Figure S4, all factors showing differential enrichment across clusters)^43^. Cluster 1 consists of primarily untranscribed elements, characterized by uniformly low TF occupancy; clusters 2–4 are composed primarily of pdTREs, and clusters 5–7 contain mostly edTREs (Figure 4B). edTREs (clusters 5-7) were preferentially enriched for pioneer factors (e.g., GATA1), and K562 lineage-specific regulators (e.g., GATA1, STAT5A, LEF1), consistent with roles in chromatin opening and cell-type-specific regulation (Figures 4C and S4-S5)^44–46^. In contrast, pdTREs (clusters 2-4) were more enriched for factors involved in core transcription processes, including Pol II subunits, general transcription factors required for preinitiation complex (PIC) assembly (TBP, TAF1, TAF9), initiation factors (NRF1, SP1, ELF4), and pausing or early elongation factors (SPT5) (Figures 4C and S4-S5)^47,48^. Notably, edTREs and pdTREs also exhibited distinct enrichment patterns for transcriptional coactivators and chromatin-remodeling complexes. edTREs showed substantially greater enrichment of the SWI/SNF chromatin-remodeling complex and the broad histone acetyltransferase EP300, which play crucial roles in generating nucleosome free regions, whereas pdTREs show higher enrichment of CHD1, which is more involved in generating properly-spaced nucleosome arrays and the EP400 complex, which can exchange H2A for H2A.Z and acetylate H4, H2A, and H2A.Z^49–51^. For coactivators that are involved in the Pol II pause-release step, including CCNT2 and BRD4 and MYC, they are enriched in both pdTREs and edTREs. Together, these findings suggest that differences in the recruitment of TFs, chromatin remodelers, and coactivators may contribute to the distinct regulatory properties of edTREs and pdTREs, potentially explaining their differential abilities to function as dTREs in the CIERA-seq system.

**Figure 4.**
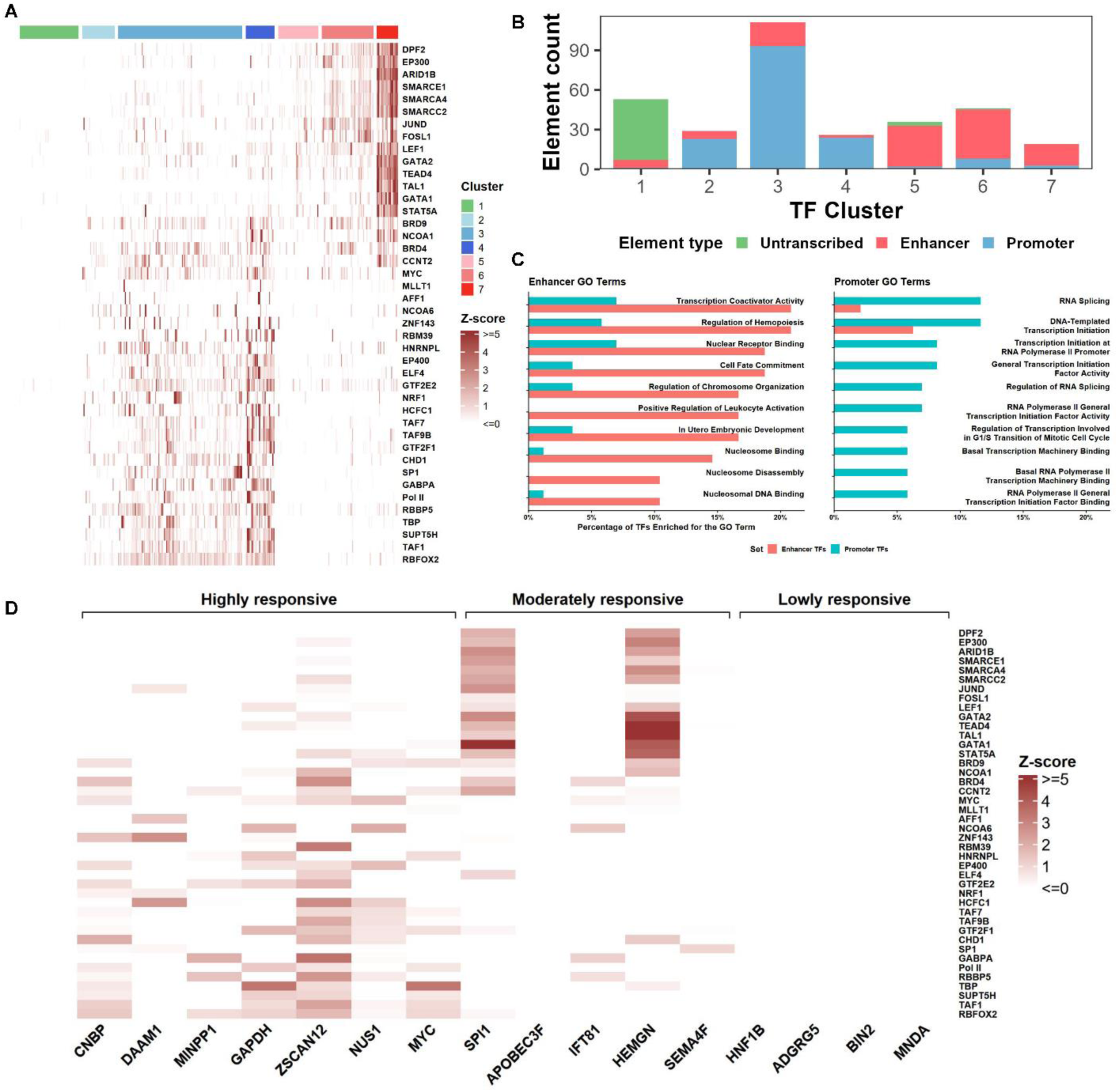
Transcription factor landscapes and functional clustering of dTREs. (A) Heatmap of ChIP–seq signals for selected transcription factors with known activator roles across dTREs. Columns represent individual dTREs, ordered by k-means clustering of transcription factor binding profiles, and rows represent transcription factors. (B) Counts of dTREs in each transcription factor–based k-means cluster. (C) Top Gene Ontology (GO) biological process (BP) and molecular function (MF) terms enriched among transcription factors preferentially enriched in edTRE-or pdTRE-dominated clusters. (D) Heatmap of ChIP–seq signals for selected transcription factors with known activator roles across the 16 target promoters.

To determine whether these patterns generalize beyond the specific dTREs included in our assay, we examined genome-wide ChIP-seq occupancy for several of the prominent implicated transcription factors across pdTREs and edTREs. This analysis recapitulated the trends observed in our selected dTRE set: enhancers showed preferential enrichment for pioneer factors and SWI/SNF chromatin remodelers, whereas promoters were more strongly enriched for general transcription factors and other factors associated with transcription initiation (Figure S3B). These genome-wide patterns suggest that the distinct TF compositions observed in our assay reflect broader differences in the regulatory architectures of promoters and enhancers rather than features unique to the dTREs selected for CIERA-seq.

Interestingly, target promoters, *CNBP*, *DAAM1*, *MINPP1*, *GAPDH*, *ZSCAN12*, *NUS1*, and *MYC*, that were highly responsive to the dTREs exhibited greater enrichment of promoter-associated transcription factors and coactivators than those that showed limited response (Figure 4D). Moderately responsive promoters, *SPI1*, *APOBEC3F*, *IFT81*, *HEMGN*, and *SEMA4F*, may depend on dTREs to supply a broader repertoire of transcriptional machinery, a requirement that fewer dTREs can satisfy, thereby limiting their activation potential. Interestingly, two moderately responsive target promoters exhibited greater enrichment of edTRE-associated transcription factors and coactivators, suggesting that the presence of these factors alone may be insufficient to support robust transcriptional activation at target promoters. Consistent with this interpretation, untranscribed target promoters, *HNF1B*, *ADGRG5*, *BIN2*, and *MNDA*, exhibited minimal enrichment of transcriptional activators and coactivators, potentially explaining their particularly low susceptibility to dTRE-mediated activation. Together, these findings suggest that the transcriptional competence of a target promoter, determined in part by its pre-existing complement of transcription factors and coactivators, may influence both the extent of its activation and the range of dTREs capable of activating it.

### Different TF clusters show distinct target promoter compatibility

We compared the target promoter-specific enhancer activities of dTREs from the seven TF-enrichment clusters using violin plots or a heatmap of activity scores across all target promoter–dTRE pairs (Figures 5A-B). Among those target promoters that are well transcribed at their native loci, consistent activity hierarchies emerged within both pdTREs and edTREs. Both types of dTREs displayed a graded increase in activity and compatibility levels from pdTRE cluster 2 to cluster 4 and edTRE cluster 5 to cluster 7 (Figures 5A-B and S6A). All active clusters exhibited substantially higher activity than cluster 1, which consists primarily of untranscribed elements. Based on these trends, we classify the clusters as inactive (cluster 1), weak, moderate and strong pdTREs (clusters 2–4), and weak, moderate and strong edTREs (clusters 5– 7).

**Figure 5.**
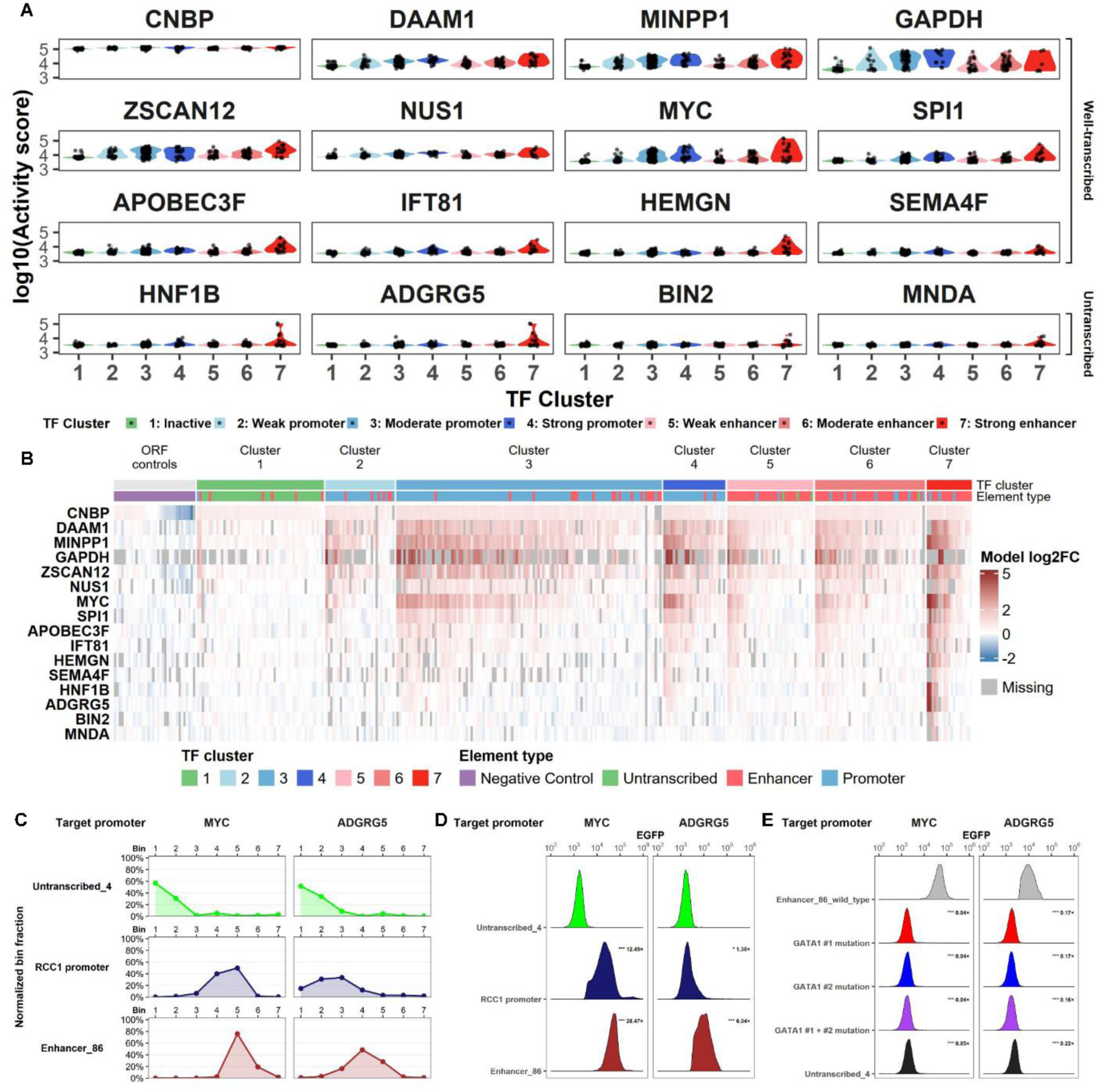
Different target promoters show different levels of activation by dTREs belonging to different TF clusters. (A) Violin plot showing the distribution of activity scores for the 16 target promoters when paired with dTREs f rom different transcription factor–based k-means clusters. (B) Log2FC of each dTRE when paired with the 16 target promoters compared to ORF derived negative controls as a heatmap. Missing promoter-dTRE pairs are shown as grey. (C) Mean normalized CIERA-seq read distributions across seven EGFP fluorescence bins for MYC and ADGRG5 target promoters paired with Untranscribed_4, the RCC1 promoter, or enhancer_86. (D) EGFP expression distributions and fold changes of cells harboring the *MYC* or *ADGRG5* target promoter paired with representative dTREs f rom different transcription factor–based clusters: untranscribed dTRE (untranscribed_4, cluster 1), strong pdTRE (*RCC1* promoter, cluster 4), and strong edTRE (enhancer_86, cluster 7). The x-axis denotes EGFP intensity. Star annotations use the Bonferroni-adjusted one-sided Welch lower confidence bound for the difference in mean log10 EGFP relative to the untranscribed control (*: bound ≥ 0.1; **: bound ≥ 0.2; ***: bound ≥ 0.5; ns: bound < 0.1). Fold change is the ratio of g eometric mean EGFP intensities. (E) EGFP expression distributions and fold changes of cells with the *MYC* or *ADGRG5* target promoter paired with the strong enhancer_86 (cluster 7), carrying different combinations of mutations in the two GATA1 motifs. Star annotations use the negative of the Bonferroni-adjusted one-sided Welch upper confidence bound for the difference in mean log10 EGFP relative to wild-type enhancer_86 (*: −upper bound ≥0.1; **: ≥0.2; ***: ≥0.5; ns: <0.1). Fold change is the ratio of geometric mean EGFP intensities.

Within both pdTREs and edTREs, strong ones showed broader and more robust enrichment of pdTRE-or edTRE-specific TFs than their moderate or weak counterparts. These functional differences were reflected in distinct transcriptional and chromatin-accessibility features at their native genomic loci, providing a mechanistic basis for the observed activity trends (Figures S6B-D)^35^. Among pdTREs, PRO-cap and H3K4me3 signals increased progressively from weak to moderate to strong pdTREs, paralleling the graded enrichment of factors involved in the core transcription process. edTREs (clusters 5–7) exhibited progressively increased chromatin accessibility, as measured by DNase-seq, consistent with their enrichment for pioneer factors and SWI/SNF chromatin remodelers. H3K27ac signal increased with dTRE strength in both pdTRE and edTRE classes. Together, these patterns suggest that enhancer strength in pdTREs is potentially driven primarily by enhanced recruitment of core transcription machinery, whereas edTREs achieve higher activity through an increased capacity to establish open chromatin via pioneer factors and SWI/SNF chromatin remodelers.

As noted above, untranscribed target promoters were activated by only a small subset of dTREs, nearly all of which belonged to cluster 7, a group of strong edTREs characterized by high enrichment of pioneer factors such as GATA1 and SWI/SNF chromatin-remodeling complexes. However, not all cluster 7 dTREs could activate these untranscribed promoters, despite exhibiting comparable enrichment of GATA1 and SWI/SNF components. To investigate the molecular features underlying this difference, we compared transcription factor and coactivator occupancy between cluster 7 dTREs that strongly activated untranscribed promoters and those with limited or no activating activity (Figure S6E). Notably, several of the most differentially enriched factors were not exclusively associated with edTREs, including GTF2F1, BRD4, ETV5, NFRKB, HMGN3, and SMAD4. These factors participate in diverse aspects of transcriptional regulation, ranging from chromatin organization and remodeling to preinitiation complex (PIC) assembly, Pol II recruitment, and promoter-proximal pause release. These observations suggest that pioneer factor recruitment and chromatin remodeling, although potentially important for establishing a permissive chromatin environment, may not be sufficient to activate transcription from otherwise untranscribed target promoters. Instead, robust activation may additionally require the recruitment of factors involved in downstream steps of the transcription cycle, including PIC assembly, Pol II recruitment, and escape of paused Pol II to productive elongation. Under this model, cluster 7 dTREs enriched primarily for pioneer factors and SWI/SNF chromatin remodelers may facilitate chromatin accessibility but many lack the full complement of transcriptional regulators necessary to support productive transcription. In contrast, dTREs of cluster 7 that are capable of recruiting both chromatin-opening factors and additional components of the transcriptional machinery overcome these limitations and activate otherwise untranscribed promoters. Together, these findings further support a transcriptional complementation model in which the ability of a dTRE to activate a target promoter depends on its capacity to provide the regulatory activities that are limiting at that promoter.

### Chromatin landscape of intrinsically closed and open target promoters when paired with promoter-derived distal TREs and enhancer-derived distal TREs

A key distinction between strong edTREs (cluster 7) and pdTREs (cluster 4) was their ability to activate untranscribed promoters. These promoters were activated almost exclusively by a subset of cluster 7 dTREs, the strongest edTRE class and the most highly enriched for pioneer factors and SWI/SNF chromatin remodelers (Figure 5A). Given the low chromatin accessibility and TF enrichment at these promoters’ native loci, we hypothesized that they remain largely inaccessible at the test locus and therefore require dTREs capable of opening chromatin for activation. To test this hypothesis, we generated cell lines in which either the *MYC* or *ADGRG5* promoter was paired with untranscribed_4 (cluster 1; inactive), the *RCC1* promoter (cluster 4; strong pdTRE), or enhancer_86 (cluster 7; strong edTRE). Consistent with the CIERA-seq bin distributions, both target promoters exhibited basal EGFP expression when paired with untranscribed_4 (Figures 5C-D). The *RCC1* promoter strongly activated *MYC* but only modestly activated *ADGRG5*, whereas enhancer_86 strongly activated both promoters (Figures 5C-D).

To assess the chromatin architectures of the target promoters in the landing-pad locus, we performed deaminase-based footprinting on promoter–dTRE constructs, focusing on *MYC* and *ADGRG5* promoters (Figure S7A). We expected that the constructs paired with an untranscribed_4 dTRE would reflect these promoter-intrinsic chromatin states in the genome-integrated context, because untranscribed_4 dTRE shows no regulatory effect on these target promoters. In this construct, the *MYC* promoter exhibited prominent accessibility, as indicated by a region with high deamination rate followed by phased downstream nucleosome pattern, whereas the *ADGRG5* promoter accessibility was indistinguishable from that of the surrounding region, and it shows no clear adjacent phased nucleosome pattern (Figure 6A). These observations indicate that the intrinsic accessibility of target promoters varies substantially. The difference in accessibility between the two promoters may, at least in part, be encoded by their underlying sequence features, as the *MYC* promoter contains a higher density of transcription factor binding motifs and shows stronger TF and cofactor ChIP-seq occupancy than the *ADGRG5* promoter (Figure 4D). We also observed pronounced differences in accessibility among the dTREs themselves. Both enhancer_86 and the *RCC1* promoter exhibited substantially higher deamination rates than the untranscribed_4 dTRE, indicating that individual dTREs possess distinct intrinsic capacities to establish accessible chromatin at the landing-pad locus (Figure 6A).

**Figure 6.**
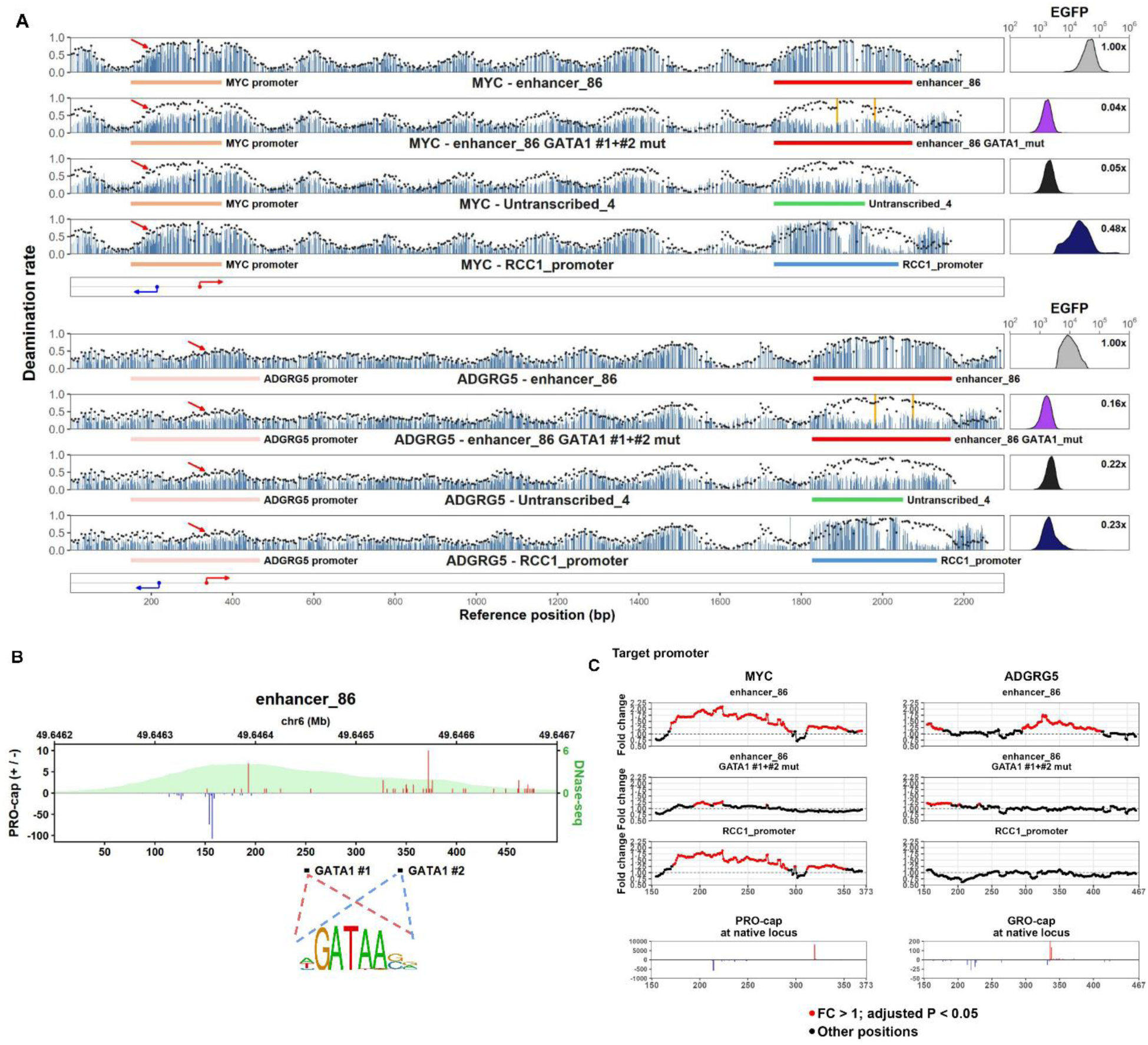
Distinct chromatin constraints underlie promoter-derived and enhancer-derived dTRE-mediated activation. (A) Average deamination rates at individual C and G positions across the indicated promoter–dTRE pairs following integration into the chromatinized landing-pad locus. The dots above each C and G in each track represent the patterns in the target promoter-enhancer_86 pair (top track) for the *MYC* and *ADGRG5* promoter series. Horizontal red and blue arrows represent PRO-cap TSS positions for *MYC* and GRO-cap TSS positions for *ADGRG5* at the corresponding endogenous promoters in cell types in which they are normally active: K562 cells for *MYC* and GM12878 cells for *ADGRG5*. Orange vertical bars represent where the TF binding motif mutations are. Plots to the right show EGFP fluorescence distributions for each promoter-dTRE combination. Fold changes are relative to the corresponding wild-type enhancer_86 pair for each target promoter. (B) PRO-cap and DNase-seq tracks of the strong enhancer_86 (cluster 7) shown in (A), highlighting the locations of two GATA 1 binding motifs and their consensus sequence. (C) Fold changes and associated P values for pooled deamination rates in 10-bp sliding windows f rom the indicated *MYC* and *ADGRG5* promoter–dTRE pairs, calculated relative to the corresponding promoter–untranscribed_4 pairs. Red dots represent FC > 1 with adjusted P < 0.05 and black for other positions. PRO-cap and GRO-cap tracks shown at the bottom indicate the positions and relative levels of transcription initiation at the corresponding endogenous promoters in cell types in which they are normall y active: K562 cells for *MYC* and GM12878 cells for *ADGRG5*. The x-axis shows positions along the *MYC* or *ADGRG5* promoter on the same scale as in (A).

Notably, our deaminase-based footprinting data indicate that, although the *MYC* promoter possesses substantial intrinsic accessibility, pairing it with either enhancer_86 or the *RCC1* promoter further increases accessibility across the promoter region (Figure 6C). This effect is particularly pronounced in the upstream region between the divergent transcription start sites, suggesting that these dTREs may promote additional recruitment or stabilization of transcription factors at the *MYC* promoter. Increased TF occupancy could in turn contribute to the formation or maintenance of a more accessible promoter architecture, thereby facilitating higher levels of transcriptional activity. In contrast, the *ADGRG5* promoter was selectively activated by enhancer_86. Consistent with this functional specificity, pairing with enhancer_86 substantially increased chromatin accessibility around the reporter-associated TSS of the otherwise poorly accessible *ADGRG5* promoter (Figure 6C). These results suggest that enhancer_86 promotes formation of a local nucleosome-depleted region at or near the site of transcription initiation, thereby creating a chromatin environment permissive for transcription from an intrinsically inaccessible promoter. Importantly, this activity of enhancer_86 depended on GATA1 binding. Mutation of the GATA1 motifs within enhancer_86 abolished the increase in enhancer accessibility, eliminated its enhancer activity when paired with ADGRG5 promoter, and prevented the associated gain in accessibility around the *ADGRG5* TSS (Figures 5E, 6A-C, and S7C). Together, these observations support a model in which GATA1-dependent activity at enhancer_86 promotes SWI/SNF dependent chromatin opening at both the enhancer and its target promoter, thereby enabling transcriptional activation of an otherwise inaccessible promoter ^52^. Notably, mutation of the GATA1 binding motifs also abolished the ability of enhancer_86 to activate the *MYC* promoter. In addition to promoting chromatin opening, GATA1 may also contribute to subsequent steps of transcriptional activation^53^.

### Role of specific TFs involved in enhancer activity of a promoter-derived distal TRE

The *RCC1* promoter displayed strong enhancer activity when paired with the *MYC* target promoter but showed very weak activity when paired with the *ADGRG5* promoter (Figures 5B and S7B). Because pdTREs are enriched for transcription factors involved in the core transcription process, we tested whether disrupting the high-confidence binding motifs for SP1, NRF1, and ELF4, which are tightly linked to TSSs and transcription initiation, would compromise their enhancer function (Figure 7A)^54–56^. We introduced individual and combinatorial mutations by fully replacing each motif with length-matched, non-regulatory sequence, and measured enhancer activity of the mutated pdTREs by pairing every mutant with the *MYC* promoter at the same genomic locus and quantifying EGFP expression (Figures 7B and S7D). Mutating a single SP1, ELF4, or NRF1 motif caused modest reductions in enhancer activity. Double mutations involving ELF4+NRF1 or SP1+ELF4 further reduced activity, whereas disruption of SP1+NRF1 led to a pronounced loss of enhancer function. Simultaneous mutation of all three motifs completely abolished enhancer activity, reducing output to levels comparable to untranscribed_4 (cluster 1; inactive). These results demonstrate that recruitment of TFs involved in initiation is required for pdTREs to function as enhancers and suggest partial functional redundancy and cooperativity among these TFs in supporting enhancer activity^56^.

**Figure 7.**
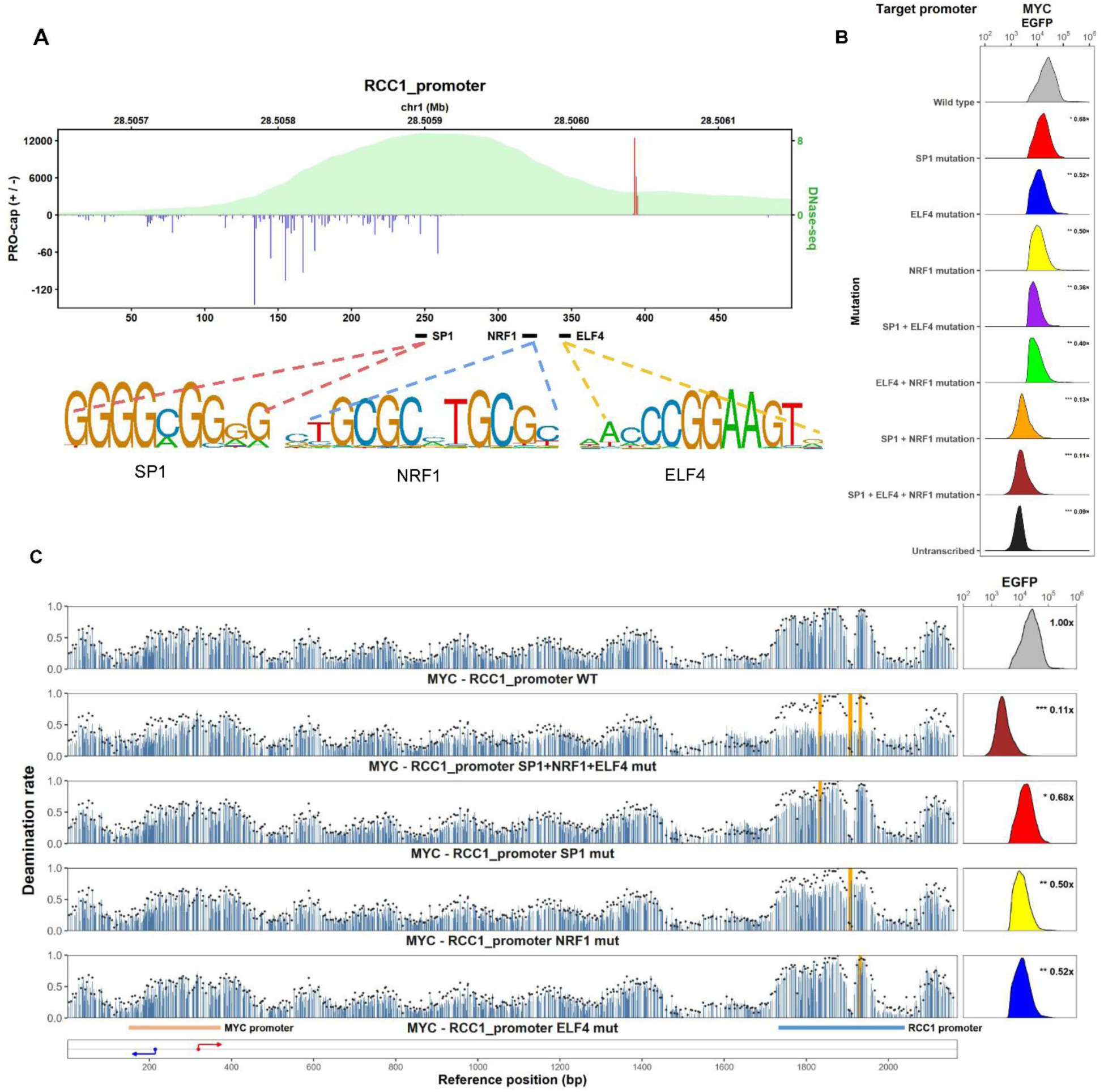
Mutations in SP1, NRF1, and ELF4 binding motifs change the RCC1 promoter’s enhancer activity and footprinting patterns. (A) PRO-cap and DNase-seq tracks of the RCC1 promoter (cluster4), highlighting the locations of SP1, NRF1, and ELF4 binding motifs and their consensus sequence. (B) EGFP expression distributions and fold changes of cells with the *MYC* target promoter paired with the *RCC1* promoter (cluster 4), carrying different combinations of mutations in the SP1, NRF1, and ELF4 binding motifs. Star annotations use the negative of the Bonferroni-adjusted one-sided Welch upper confidence bound for the difference in mean log10 EGFP relative to wild type (*: −upper bound ≥ 0.1; **: ≥ 0.2; ***: ≥ 0.5; ns: < 0.1). Fold change is the ratio of geometric mean EGFP intensities. (C) Average deamination rates at individual C and G positions across the indicated promoter–dTRE pairs following integration into the chromatinized landing-pad locus. Orange vertical bars indicate mutated TF motifs; black dots show the wild-type reference accessibility pattern. The EGFP profiles on the right show cell fluorescence distributions; fold changes and stars are defined in B.

We further analyzed several of these *MYC*-*RCC1* promoter mutant lines using deaminase-based footprinting and identified additional TF-dependent patterns of chromatin accessibility. At the *RCC1* promoter, we observed several localized protection footprints, one of which appears to be associated with NRF1 binding, as mutation of the NRF1 motif specifically eliminated this footprint (Figure 7C). However, disruption of NRF1 binding alone had only a modest effect on the enhancer activity of the *RCC1* promoter and did not affect paired *MYC* promoter accessibility, suggesting that other transcription factors may compensate for the loss of NRF1 function (Figures 7B-C). Similarly, individual mutations of the SP1 or ELF4 motifs produced only modest changes in accessibility at either the *RCC1* promoter or the paired *MYC* target promoter (Figure 7C). In contrast, simultaneous mutation of the NRF1, SP1, and ELF4 motifs resulted in a near-complete loss of the nucleosome-free region at the *RCC1* promoter, accompanied by a corresponding loss of its enhancer activity (Figure 7C). These results suggest that NRF1, SP1, and ELF4 act cooperatively to establish or maintain accessible chromatin at the *RCC1* promoter, potentially in a SWI/SNF independent manner^57,58^. The loss of *RCC1* promoter-mediated transcriptional activation was accompanied by only a slightly reduced accessibility at the *MYC* target promoter. These findings suggest that the loss of enhancer activity may involve defects beyond simply reduced target promoter accessibility. NRF1, SP1, and ELF4 may therefore contribute to multiple steps in transcriptional regulation downstream of the chromatin opening step, potentially including transcription initiation and the release of paused Pol II into productive elongation.

## DISCUSSION

In this study, we developed a chromatin-integrated landing pad–based assay, CIERA-seq, to systematically interrogate regulatory effects on gene expression from the combinatorial interactions between ∼350 dTREs and 16 target promoters in a native chromatin context^25^. Unlike prior plasmid-based approaches that focused primarily on enhancer–promoter interactions^6–10^, we explored both enhancer–promoter and promoter–promoter regulatory interactions. Importantly, we find that the chromatin structures of particular CIERA-seq integrated promoter-dTRE combinations can be effectively examined by a deaminase-based footprinting approach. Comparison of the footprints on CIERA-seq constructs with different dTRE combinations revealed coordinated changes in promoter and dTRE accessibility, as well as the effects of mutations in key TF-binding motifs. The combined analysis of gene activity and chromatin structure along continuous DNA fibers spanning the entire promoter-dTRE reporter constructs provided a powerful means of testing general principles of how dTRE types cooperate with different promoter types to regulate transcription and associated chromatin structure.

Consistent with the model in which promoter activity relies on cooperation with surrounding dTREs, we find that when promoters are removed from their native chromatin environments and placed into a standardized landing pad with minimal flanking dTREs, their transcription output frequently decreases. Promoters with strong transcription at their native loci sometimes show, not surprisingly, markedly attenuated activity in isolation, underscoring that local TRE and chromatin context exerts dominant control. Using our CIERA-seq system, we find that target promoters can be stimulated not only by canonical enhancers but also by promoters. This extends recent large-scale plasmid-based evidence that promoters can adopt enhancer-like roles and suggests that TRE–TRE crosstalk is a general feature of transcription regulation^6^.

Pol II must progress through multiple distinct steps of the transcription cycle to produce mRNA. At many of these steps, TFs and their associated coactivators modulate the progress of Pol II. The steps involve the opening of chromatin structure by pioneer TFs and SWI/SNF chromatin remodelers, PIC assembly, Pol II initiation, early elongation and promoter proximal pausing, and the maturation and release of Pol II into a productive elongation complex^59^. Some promoters are limited at the earliest step due to an inability to recruit chromatin-opening factors, whereas others can be limited at later stages such as PIC assembly or pause release^60^. Different TFs and coactivators are known to bind a promoter and their complementary activities can cooperate to drive transcription^39,45,61,62^. Likewise, it’s easy to imagine that entire TREs, both promoters and enhancers, could also collaborate through providing complementary TFs and coregulators to drive high levels of transcription activity.

Our results suggest that pdTREs and edTREs contribute to distinct regulatory activities that reflect differences in the transcriptional machinery they recruit. Across the dTREs examined, edTREs were preferentially enriched for pioneer factors, SWI/SNF chromatin remodelers, and lineage-specific regulators, and specific coactivators including EP300, whereas pdTREs were more strongly enriched for Pol II, general transcription factors, specific coactivators like EP400, and factors associated with transcription initiation, pausing, and early elongation; importantly, similar distinctions were observed genome-wide. Despite these differences, both classes readily can act as enhancers in our assay when paired with compatible target promoters. For target promoters that are intrinsically accessible, enhancer potencies of edTREs or pdTREs tend to scale with the overall degree of binding of transcription factors. Compositional differences of dTREs correlated with distinct patterns of target-promoter compatibility. Target promoters with greater pre-existing enrichment of transcription factors tend to be activated by a broader range of dTREs and generally exhibit stronger activation by pdTREs than by edTREs, whereas a small subset of strong edTREs showing high enrichment of pioneer factors and SWI/SNF remodelers displayed broader compatibility and were uniquely capable of activating transcriptionally inactive promoters with low TF occupancy and accessibility. Together, these observations argue that dTRE strength cannot be described by a single universal activity scale; instead, regulatory output depends on whether the molecular activities provided by a dTRE complement or simply add to the regulatory state and requirements of its target promoter.

Our chromatin-accessibility and mutational analyses provide a mechanistic insight for this compatibility. Deaminase-based footprinting revealed that the *MYC* promoter maintains substantial intrinsic accessibility and clear downstream nucleosome phasing patterns after integration at the safe harbor locus. On the other hand, the untranscribed *ADGRG5* promoter remains largely inaccessible with no clear downstream nucleosome phasing patterns. Previous work has shown that activation of promoters with low accessibility and poorly phased adjacent nucleosomes requires SWI/SNF chromatin remodelers ^63^. This may help explain why the *ADGRG5* promoter is activated only by the strong edTRE enhancer_86 enriched with SWI/SNF binding, whereas the *MYC* promoter can be activated by both enhancer_86 and the *RCC1* promoter. GATA1 motif disruption abolished enhancer_86-dependent chromatin opening and activation of *ADGRG5*, supporting a role for edTRE-enriched pioneer factors and associated SWI/SNF in overcoming chromatin barriers; however, GATA1 was also required for activation of the already accessible *MYC* promoter, indicating that its contribution extends beyond chromatin opening alone^53^. Conversely, at the *RCC1* promoter-derived dTRE, combined disruption of SP1, NRF1, and ELF4 eliminated enhancer activity with only minor effects on the openness of the MYC promoter, consistent with promoter-enriched initiation factors contributing cooperatively to steps downstream of chromatin opening during their distal transcription activation. These findings therefore support a model in which different dTREs provide partially distinct regulatory functions—from establishing accessibility to supplying or stabilizing transcriptional machinery— and target promoters respond according to which of these functions are limiting in their existing regulatory state.

### Limitations

A couple of limitations of this study should be noted. First, TF occupancy was inferred from ChIP-seq signals measured at the endogenous loci of dTREs and used to classify dTREs into distinct clusters. However, TF binding at these native loci may not fully recapitulate occupancy following relocation of dTREs to the landing pad context. Despite this caveat, for many TFs with well-defined DNA-binding motifs, occupancy is expected to be largely sequence-driven and therefore broadly preserved across genomic contexts. Second, our experimental framework for evaluating transcriptional regulation only captures chromatin opening, some TF binding, and overall transcriptional output. This approach does not distinguish between intermediate regulatory processes, such as PIC assembly, promoter-proximal pausing, or pause release. Dissecting the contributions of individual dTREs to these mechanistic sub-steps will require the development of more refined experimental systems with higher temporal and mechanistic resolution. Third, CIERA-seq is designed to systematically characterize the regulatory potential of dTREs and the differential responsiveness of promoters under defined conditions of relative proximity. Therefore, regulatory interactions observed between dTREs and promoters in CIERA-seq do not necessarily indicate that these elements interact or regulate one another in much more distal positions as may be present at their endogenous genomic context.

## RESOURCE AVAILABILITY

### Lead Contact

Further information and requests for resources and reagents should be directed to and will be fulfilled by the lead contact, John Lis.

### Materials Availability

Plasmids generated for this study are available upon request and will be fulfilled by the lead contact, John Lis.

### Data and Code Availability

All raw and processed data of the CIERA-seq and long-read deamination sequencing can be found at NCBI’s Gene Expression Omnibus under accession number GSE316493.

## ACKNOWLEDGEMENTS

This work was supported by the National Institutes of Health grant R01 HG012970 to J.T.L. and H.Y.

## AUTHOR CONTRIBUTIONS

Y.J. conceptualized the study with guidance from Y.H., Z.Z., H.Y, and J.T.L. Y.J. generated the CIERA-seq data with the help from J.L. Y.H. performed the deamination-based footprinting experiments. Z.Z. generated the landing pad containing cell line. A.K-Y.L selected the target promoters tested.

## SUPPLEMENTAL INFORMATION

Figures S1-S7, Tables 1-3.

## METHODS

### Genome build

All genomic analyses were performed using the human genome assembly GRCh38/hg38 as the reference.

### Construction of Bxb1 recombination vectors

The donor vector for Bxb1-mediated recombination was derived from a modified eSTARR-seq plasmid and the attB1-EF1a-EGFP-attB2 construct described previously^11,26^. The eSTARR-seq backbone was digested with AflIII (NEB, R0541S) and MfeI (NEB, R3589S) to remove the luciferase reporter and the SV40 polyadenylation signal. In parallel, the EGFP–polyA cassette was PCR-amplified from attB1-EF1a-EGFP-attB2 using primers designed to introduce homology arms compatible with the digested backbone. The forward primer annealed upstream of EGFP and provided homology to the 5’ end of the cleaved vector. The reverse primer annealed downstream of the SV40 polyA region and introduced a unique 9-bp randomized barcode, enabling unambiguous identification of each target promoter during sequencing. The PCR product was assembled into the digested backbone using Gibson Assembly® Master Mix (NEB, E2611), and the resulting constructs were transformed into One Shot™ ccdB Survival™ 2 T1R competent cells (Invitrogen, A10460). Transformants were selected on LB agar containing ampicillin and chloramphenicol. Sixteen colonies were picked, plasmid-prepped, and Sanger-sequenced to confirm the presence of distinct 9-bp barcodes lacking transcription-factor motifs or canonical regulatory elements. These barcoded plasmids were subsequently digested with AvrII (NEB, R0174S) and AseI (NEB, R0526S) to allow cloning of sixteen different target promoters.

### Target promoter cloning

Target promoters (sequences available in Supplementary Table 2 (target promoter info)) were PCR-amplified from K562 genomic DNA using PrimeSTAR® GXL DNA Polymerase (TaKaRa, R050B) for 32 cycles with primer pairs engineered to introduce AvrII (CCTAGG) and AseI (ATTAAT) restriction sites, with the order of sites determined by whether the primary transcript of the promoter lies in the sense or antisense orientation (primer sequences are provided in Supplementary Table 3). Promoter fragments were designed to extend at least 60 bp downstream of the divergent transcription start site to encompass promoter - proximal pausing regions. PCR products were digested with AvrII (NEB, R0174S) and AseI (NEB, R0526S) at 37 °C for 1 h, resolved by electrophoresis on a 1% agarose gel, and purified using a QIAquick Gel Extraction Kit (QIAGEN, 28704). Purified inserts were ligated into barcode-indexed donor vectors (described above) using T4 DNA Ligase (NEB, M0202S) at 16 °C overnight, transformed into the same competent cells as before, plated on the same selective plates as before, and plasmids were recovered using a QIAprep Spin Miniprep Kit (QIAGEN, 27104).

### dTRE library selection and cloning

We selected dTREs from the sets defined and tested by Tippens *et al.* and Paramo *et al.* using ChromHMM and GRO-cap^6,11^. In total, 354 dTREs were chosen, including 116 edTREs, 154 pdTREs, 50 untranscribed elements, and 34 open reading frame–based negative controls (see Supplementary Table 1 (dTRE info) for sequences and genomic coordinates). The selected genomic spans of the 320 non-ORF dTREs ranged from 152 to 553 bp (mean, 320 bp). Among the 270 pdTREs and edTREs, 200 had genomic spans of 200– 450 bp (mean across all 270, 338 bp). They were only tested in the forward genomic orientation since enhancers show function in an orientation-independent manner. All dTREs were previously cloned using a boundary extending ∼60 bp from the divergent PRO-cap TSSs to not only include the core promoter region, but also the promoter-proximal pausing region. For each target promoter, the dTRE pool was transferred into the corresponding attB1–promoter–EGFP–barcode–attB2 backbone via LR recombination. Briefly, LR reactions contained 10 µL of the pooled entry plasmids (100 ng µL⁻¹), 10 µL TE buffer, 10 µL of the matched attB1–target-promoter–EGFP–barcode–attB2 plasmid (100 ng µL⁻¹), and 10 µL LR Clonase, and were incubated at 16 °C overnight. Reaction products were transformed into E. coli TOP10 cells (5 µL LR reaction per 50 µL cells) and plated on LB–ampicillin agar plates. Colonies were collected by scraping (Avantor, 734-2602), and plasmids were purified using the E.Z.N.A.® Endo-Free Plasmid DNA Maxi Kit (Omega, D6926-00S). The resulting constructs constitute the attB1–target-promoter–EGFP–barcode–dTRE–attB2 library.

### Genomic integration of target promoter–dTRE constructs

For each of the 16 target promoters, 10 µg of the corresponding attB1–target-promoter–EGFP–barcode– dTRE–attB2 plasmid was mixed with 10 µg of pFL9_pCAG-NLS-HA-Bxb1 (Addgene 51271). DNA mixtures were concentrated by speed vacuum to ∼10 µL to maximize electroporation efficiency. Constructs were delivered into K562 landing-pad cells using the Cell Line Nucleofector® Kit V (Lonza, VCA-1003) and a Lonza Nucleofector 2b device, electroporating 10 million cells per reaction (tenfold above the manufacturer’s recommended cell number). Following electroporation, cells were recovered for 10 min in outgrowth medium composed of 70% fresh RPMI 1640 + GlutaMAX (Gibco, 61870036), 10% FBS (Avantor, CNS89510-194), and penicillin–streptomycin (Corning, 30-001-CI) mixed with 30% conditioned growth medium of identical composition. Cells were plated into 10-cm dishes and cultured for 1 week. Successfully recombined cells were isolated by fluorescence-activated cell sorting (Sony MA900), gating for loss of BFP expression from the landing-pad cassette. Integration efficiencies of ∼10% were routinely obtained. To maintain representation of the dTRE library for each promoter, at least 5 × 10⁵ BFP-negative cells were collected. Sorted cells were expanded for an additional two weeks to allow degradation of residual plasmid DNA prior to downstream assays. For those cell lines that contain one specific promoter-dTRE pair, they are also integrated using this strategy, and no single colony is selected to prevent single colony effects.

### Sorting cells into EGFP-expression bins

Following three weeks of culture in RPMI supplemented with 10% FBS and penicillin–streptomycin, cells harboring target promoter–dTRE integrations were pooled (5 × 10⁶ cells from each promoter pool; 0.8 × 10⁸ cells total), washed twice with 100 mL ice-cold PBS (Gibco, 10010023), resuspended in 10 mL, and filtered through Falcon® round-bottom tubes with 35-µm cell-strainer caps (Corning, 352235). Cells were sorted on a Sony MA900 cell sorter based on EGFP fluorescence intensity. Seven gates were drawn to correspond to defined percentiles of the EGFP expression distribution, and approximately 5 × 10⁵ cells were collected per bin. Genomic DNA from each sorted population was isolated using the DNeasy Blood & Tissue Kit (QIAGEN, 69504) and stored for downstream library preparation.

### Library preparation from genomic DNA of EGFP-expression bins

Genomic DNA from each fluorescence-sorted bin was used as template for PCR amplification of cassette fragments containing both the target-promoter barcode and the integrated dTRE sequence. A forward primer binding upstream of the promoter barcode (seq2-EGFP-barcode-2-F) and a reverse primer annealing downstream of the genomic Bxb1 integration junction on the genomic DNA but not on plasmids (seq1-ZZ103-R) were used to amplify the integrated cassette for 10 cycles (25 µL Q5® High-Fidelity 2X Master Mix, NEB M0492S; 1 µg genomic DNA; 1 µL 10 µM of each primer; H₂O to 50 µL total volume). All genomic DNA recovered from each bin was used to ensure sufficient library complexity. Following amplification, reactions were treated with Exonuclease I (NEB M0293S; 50 µL PCR product + 6 µL ExoI buffer + 3 µL H₂O + 1 µL ExoI) to remove residual primers, then purified with the DNA Clean & Concentrator-5 kit (Zymo, D4014) and eluted in 20 µL H₂O. Purified products were subsequently re-amplified for 15 cycles to append Illumina P5 and P7 adapters using primers P5-seq1 and P7-barcode-seq2. Final libraries were resolved on a 6% TBE polyacrylamide gel, and the band corresponding to the expected size was excised, crushed, and eluted overnight in TE buffer at 4 °C. Gel eluates were filtered through Spin-X 0.45-µm centrifuge tubes (Costar, 8163) and purified by phenol–chloroform extraction, then eluted in 20 µL H₂O. Library concentrations were quantified by Qubit, and equal mass from each expression bin was pooled and submitted for 2 × 150 bp paired-end sequencing (Novogene).

### Sequencing alignment and computational analysis

Adaptor trimming was performed using fastp, yielding paired outputs: Read 2 sequences containing the 9 - bp barcodes that encode target promoter identity, and Read 1 sequences corresponding to dTRE inserts. Each read set was aligned with Bowtie2 to custom reference FASTA files consisting solely of the expected promoter barcodes or dTRE sequences, respectively. For every paired-end read, the mapped promoter barcode and dTRE identifier were joined and tallied to generate counts of each target promoter–dTRE combination within each bin. For each bin, raw counts for all promoter–dTRE pairs were summed to obtain the total number of aligned reads. The abundance of each pair was expressed as a fraction of this bin total. Fractions across all seven bins were normalized such that values for each target promoter –dTRE pair summed to one, yielding a distribution that reflects its relative representation across the EGFP bins (Fig ure S1A). Because the fluorescence distribution of each promoter–TRE pair is expected to be approximately normal on the EGFP scale, reads assigned to bins that deviate substantially from this distribution may reflect technical errors during sorting. To minimize the influence of such outliers, we fitted a normal distribution to the bin-wise fluorescence profile and selected the four consecutive bins with the highest cumulative probability under the fitted distribution. The regulatory activity score was then calculated as the weighted mean fluorescence across these four bins, using the observed fraction of reads in each bin as weights, normalized by the total read fraction within the selected window (Figure S1A). It is important to note that basal EGFP expression and the limited detection range of the reporter assay may compress the apparent differences in activation scores between strong and weak dTREs, resulting in smaller observed differences than those measured by qPCR-based assays.

Three independent biological replicates were performed for each promoter–dTRE pair. Following read-count filtering, an additional replicate-level quality-control step was applied to exclude outliers. Individual replicate scores deviating from the mean across replicates by more than three times the promoter-specific standard deviation estimated using an intercept-only limma model fitted separately for each promoter were excluded. Promoter–dTRE pairs were retained for downstream analysis if at least two biological replicates passed this filtering. For significance testing, each retained replicate score was expressed as a log2 ratio relative to the arithmetic mean activity of quality-controlled ORF-derived negative controls for the same promoter and biological replicate. These control-normalized scores were analyzed using an intercept-only limma model fitted separately for each promoter, with robust, abundance-dependent empirical Bayes moderation to stabilize variance estimates. A two-sided moderated t-test was used to assess whether the mean log2 ratio differed from zero, and P values were adjusted globally using the Benjamini–Hochberg procedure. TREs were considered significantly activating if the adjusted P value was below 0.05 and the estimated log2 fold change was at least 0.6.

### K-means clustering of dTREs based on TF ChIP-seq signals

ENCODE ChIP-seq datasets obtained through the UCSC Genome Browser provided TF occupancy measurements at the tested dTREs^35,41,42^. The final grouping used 153 selected transcription factors and cofactors across 320 genomic dTREs; the 34 ORF controls were not TF-clustered. Initial k-means clustering was followed by subdivision, reassignment, and merging to obtain seven fixed clusters containing 53, 29, 111, 26, 36, 46, and 19 dTREs. Per-factor z-scores for the dTRE heatmaps were calculated across all 320 genomic dTREs using the sample standard deviation. The displayed color scale saturates values at or below 0 and at or above 5. The target-promoter heatmap used the joint reference of 320 dTREs and 16 target promoters.

### Metaplot analysis of chromatin and transcription features

Metaplots of chromatin accessibility and histone modification (DNase-seq (GSM816655), H3K27ac (GSM733656), H3K4me3 (GSM788087), and H3K4me1 (GSM788085)), nascent transcription (PRO-cap and PRO-seq), and selected transcription factor ChIP–seq datasets were generated in R using the GenomicRanges, IRanges, rtracklayer, tibble, dplyr, tidyr, and ggplot2 packages. For all datasets except PRO-seq, signal was extracted in a ±500 bp window centered on each dTRE to capture proximal regulatory activity. For PRO-seq, a ±5 kb window was used to quantify transcription output across the gene body. Group-mean profiles were normalized by a common maximum absolute group-mean value for each track across the compared categories, enabling comparison across datasets on a standardized 0–1 scale. ChIP– seq tracks for individual transcription factors were downloaded from ENCODE and processed using the same ±500 bp genomic boundaries. For the genome-wide analysis in Figure S3B, 43,927 hg38 regions were classified using direct overlap with annotated protein-coding or lncRNA transcript TSSs from GENCODE v49: 20,107 promoter-like regions and 23,820 enhancer-like regions. Signals were summarized in 100 spatial bins spanning ±500 bp from each region center without strand flipping. For each factor, promoter and enhancer mean profiles shared a normalization denominator equal to the larger group-mean maximum. Within each class, heatmap rows followed descending RNA polymerase II ChIP-seq signal.

### Gene ontology analysis of transcription factors enriched at pdTREs and edTREs

Gene ontology analysis of transcription factors differentially enriched between pdTREs and edTREs was performed using clusterProfiler, org.Hs.eg.db, and enrichplot in R, with a false-discovery rate threshold of q ≤ 0.05. Representative Gene Ontology terms in the Biological Process and Molecular Function categories were selected for visualization. Network representations were generated using ggraph, ggplot2, and grid, with nodes corresponding to GO terms and transcription factors contributing to each term.

### Flow-cytometry quantification

For flow-cytometry panels, up to 8,000 events per sample were sampled from the exported event files using random seed 42; nonpositive or nonfinite EGFP values were excluded. Fold change was calculated as 10 raised to the difference in mean log10 EGFP between sample and reference, equivalent to a ratio of geometric means. One-sided Welch confidence bounds used a Bonferroni-adjusted alpha of 0.05 divided by the number of comparisons within each promoter and figure family. Positive-effect stars used lower bounds of at least 0.1, 0.2, or 0.5 log10 units; negative-effect stars used upper bounds of at most −0.1, −0.2, or −0.5. These cell-level annotations describe the sampled fluorescence distributions and do not establish between-experiment biological reproducibility. Boxplots show medians and interquartile ranges, with whiskers extending to 1.5 times the interquartile range.

### Deamination-based footprinting for DNA accessibility at the landing pad locus

To minimize variation in deamination reaction conditions across samples, we pooled K562 cells containing different specific promoter-dTRE pairs to test together to 2-4 M cells per experiment to start with. Cells were first washed two times with PBS on ice. They were then permeabilized by incubating them with 1.5 ml of Permeabilization buffer (10 mM Tris-HCl (pH 7.4), 300 mM Sucrose, 10 mM KCl, 5 mM MgCl2, 1 mM EGTA, 0.1% NP-40, 0.05% Tween-20, 0.5 mM DTT, 1x Protease Inhibitor (Thermo Fisher, A32965), and 4 U/ml SUPERase-In RNase Inhibitor in nuclease-free water) on ice for 5 min. The nuclei were washed twice with 1 ml of the same buffer. The nuclei were then prewashed once with ice-cold deamination buffer (50 mM Bis-Tris (pH 6.0), 0.1% Triton X-100 in nuclease-free water) and resuspended in the same buffer at room temperature. The nuclei were counted and adjusted to 1×10^6^ nuclei/ml using the deamination buffer, and 200 ul of the nuclei were mixed with 20 ul of in vitro translated CseDa01 in PURExpress (NEB, E6800S) and incubated in a 37 °C heat block for 18 minutes to convert accessible cytosines in chromatin to uracil^64^. The reaction was then quenched with 12 ul of 10% SDS and treated with 1 ul of Proteinase K (Invitrogen, AM2546) for 1 hour at 56 °C, followed by phenol-chloroform extraction and isopropanol precipitation. The purified genomic DNA was treated with 1 ul of RNase A (Thermo Fisher, EN0531) at 37 °C for 1 hour, followed by a final purification with Genomic DNA Clean & Concentrator-10 (Zymo Research, D4011). DNA concentrations were measured by both Nanodrop and Qubit dsDNA HS kit (Thermo Fisher, Q33231). Once genomic DNA was extracted, PCR was performed to amplify the genome-integrated reporter region on both the top and bottom strands using LongAmp Taq 2X Master Mix (NEB, M0533S), using two specific sets of primers designed for regions containing up to two cytosines. At the positions corresponding to cytosines, the primer nucleotide was either C or T. The resulting libraries were run on an agarose gel and gel-extracted, followed by Nanopore sequencing by Plasmidsaurus.

### Nanopore sequencing data processing

Reads were aligned using minimap2 to the reference sequences in a FASTA file corresponding to the reporter constructs in the pooled samples. We then separated the alignments into top-strand-derived and bottom-strand-derived groups based on C-to-T and G-to-A conversions per alignment using a custom script. The two strand-derived groups were processed separately until the downstream step described below. To minimize misalignments, we retained only primary alignments longer than 2 kb, which covered nearly the entire genome-integrated reporter construct. For constructs carrying mutations in TF-binding motifs, we further verified the mutation patterns in these filtered alignments using a custom script that accounted for C-to-T conversion. Deaminase-induced C-to-T conversions and G-to-A conversions observed on the top-strand reference were used to infer accessible regions on the top and bottom DNA strands, respectively. The deamination rates were calculated across individual fibers at each C and G position, and the C-to-T (top-strand) and G-to-A (bottom-strand) measurements were then combined into a single bar-plot track for each sample for direct comparison of chromatin accessibility patterns across promoter–dTRE pairs. For deamination profiling, top-strand C-to-T and bottom-strand G-to-A changes were counted relative to each construct’s matching reference, including the designed mutant sequence. Unmapped, non-primary, and QC-failed alignments were excluded, together with reads showing more than five opposite-strand conversions or failing construct-identity mutation checks at diagnostic sites where mutations are introduced. At each eligible position, the rate was the number of reads showing C-to-T or G-to-A changes, respectively, divided by the total coverage at that position. Top-and bottom-strand rates were displayed together along the construct reference, alongside matched EGFP distributions and transcription-initiation tracks.

### CRISPRi data analysis to identify regulatory effects of dTRE silencing

CRISPR interference screening data from Gasperini *et al.*^24^ were reanalyzed to evaluate the effects of promoter and enhancer perturbation on expression of neighboring genes ^24^. We excluded gRNA groups targeting promoters of genes with transcription-related Gene Ontology (GO) annotations to minimize potential trans effects. The plotted analysis was restricted to tested perturbation–gene pairs with decreased expression. Significant repression was defined as a transcript fold change <1 and an empirical adjusted P value <0.10, the same cutoffs used in the original paper. Promoter–promoter analyses excluded self-targeting controls and pairs with transcription start sites separated by ≤1 kb. To incorporate chromatin contacts, targeted-region and responding-gene TSS coordinates were converted from hg19 to hg38 and intersected with Micro-C loop anchors expanded by 20 kb on each side. A pair was classified as having contact when its two endpoints overlapped opposite anchors of the same loop, allowing either orientation. Contact status was used to stratify the results. Within each interaction class, the fraction of significantly repressed pairs was calculated separately for contact and no-contact groups among pairs showing decreased expression.

## SUPPLEMENTAL FIGURES

**Supplemental Figure 1.**
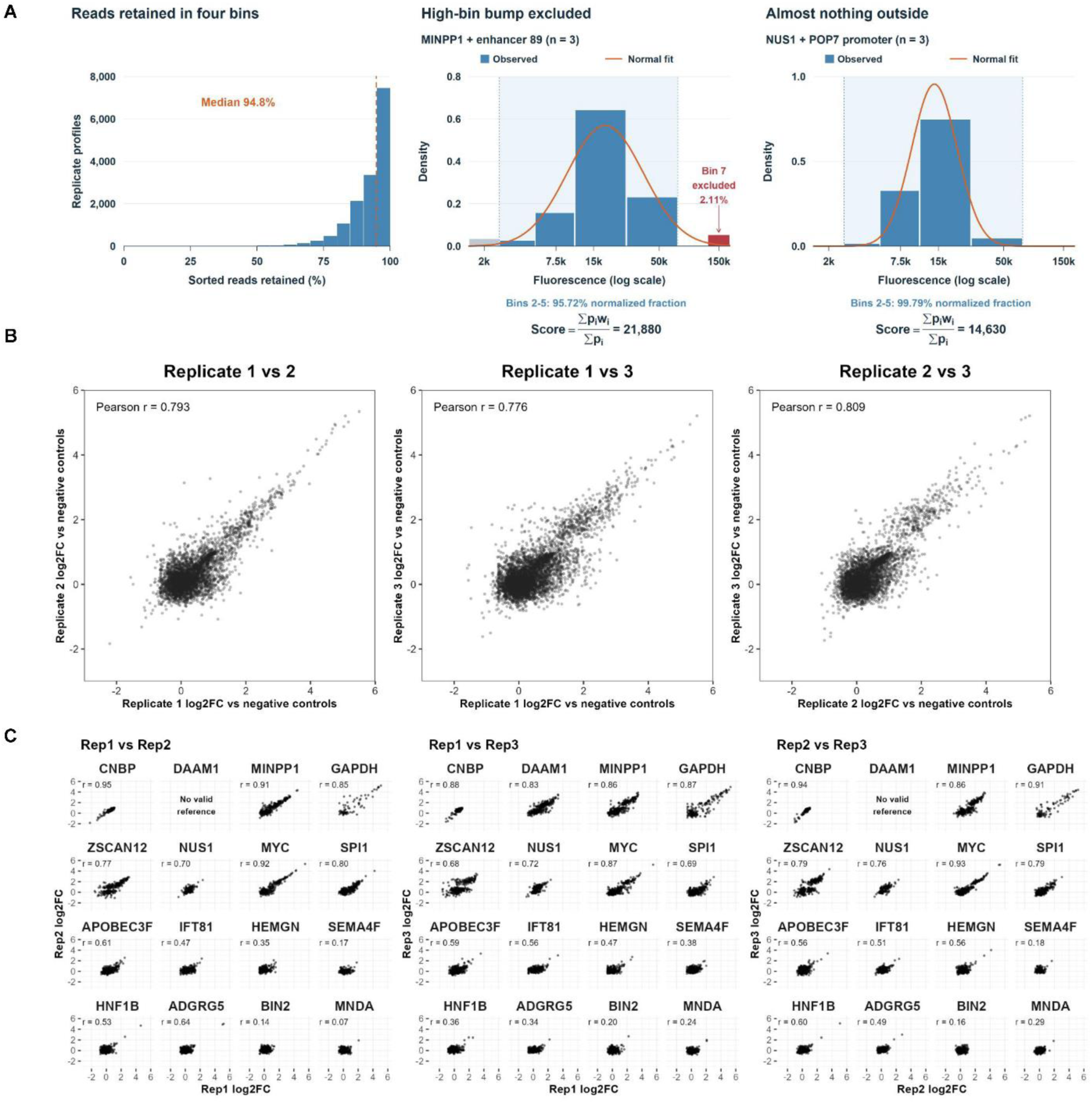
Activity scoring and reproducibility of CIERA-seq. (A) Left, percentage of sorted reads retained within the selected four-bin window across replicates. The four-bin window is defined as the four consecutive EGFP fluorescence bins with the highest cumulative probability under the fitted normal distribution. Right, representative examples illustrating activity score calculation, including exclusion of a high-fluorescence outlier peak (MINPP1– enhancer 89) and near-complete read retention (NUS1–POP7 promoter). Orange curves represent normal distributions fitted on the log-fluorescence scale; blue bars indicate selected bins, while gray and red bars indicate excluded bins. Activity scores were calculated as weighted mean fluorescence values using observed read fractions normalized within the selected four-bin window. (B) Correlation of log2FC of activity scores against ORF negative controls f rom all promoter-TRE pairs of the three biological replicates. (C) Correlation of log2FC of activity scores against ORF negative controls for each promoter between pairs of the three biological replicates. Reduced correlation between biological replicates for certain promoters was predominantly attributable to their weak responsiveness to dTREs, which yielded low dynamic range in activity scores. DAAM1 rep2 had low read counts and was excluded.

**Supplemental Figure 2.**
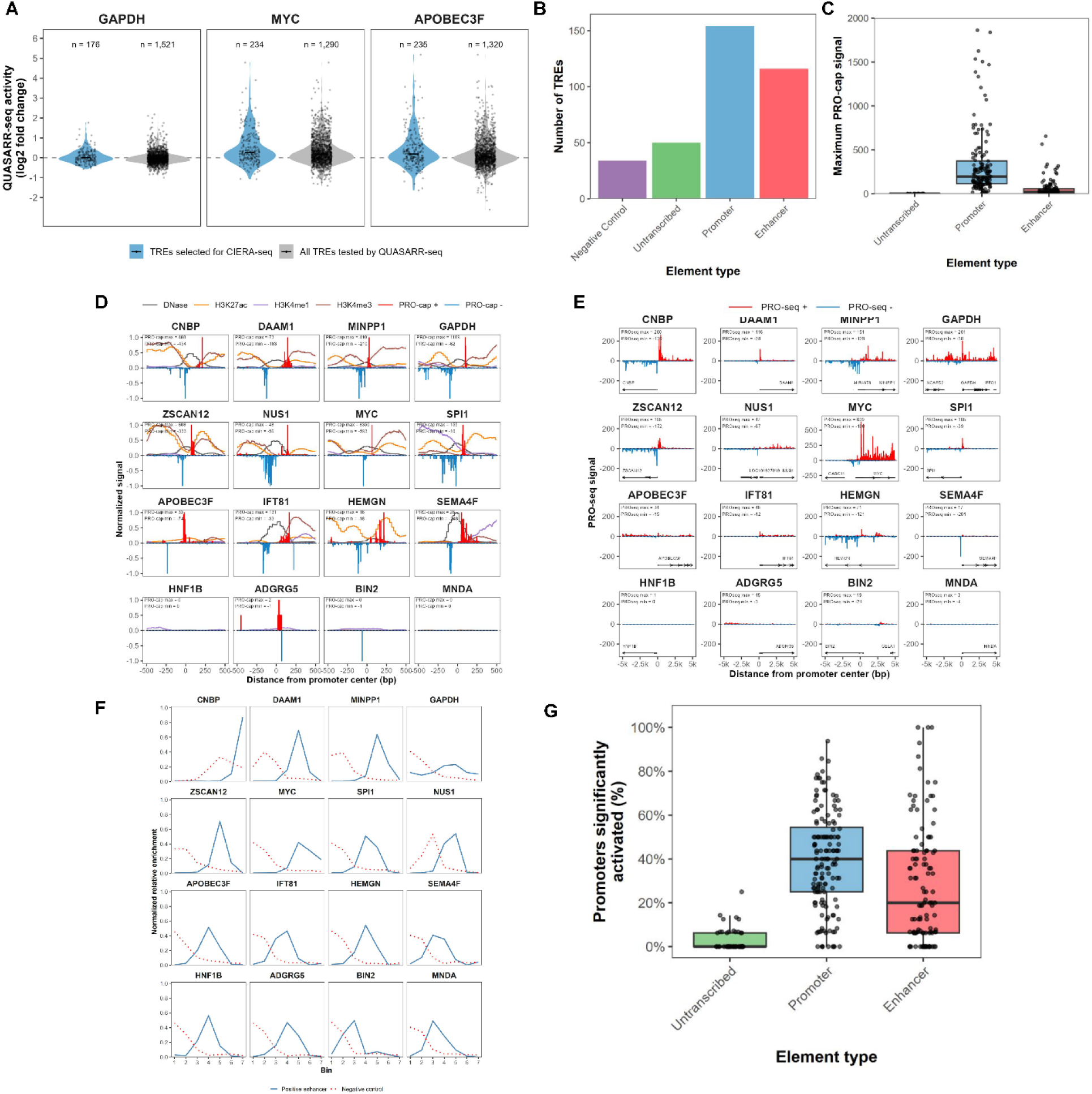
Intrinsic activity, chromatin and transcriptional profiles of target promoters, and comparison of differences in features and activity of promoter-derived dTREs and enhancer-derived dTREs. (A) Distributions of QUASARR-seq activity (log2 fold change) for TREs selected and well cloned for CIERA-seq (blue) and all TREs tested by QUASARR-seq (gray), shown for the GAPDH, MYC, and APOBEC3F target promoters. (B) Counts of dTREs classified by regulatory nature. (C) Boxplots of maximum PRO-cap signal across dTREs at native loci of different regulatory natures. (D) Profiles of PRO-cap, DNase-seq, and ChIP–seq signals for H3K27ac, H3K4me1, and H3K4me3 across the 16 target promoters at their native genomic loci, shown f rom −500 bp to +500 bp relative to the promoter center. (E) PRO-seq profiles across the 16 target promoters, shown f rom −5 kb to + 5 kb relative to the promoter center. (F) Normalized relative enrichment distributions of the 16 promoters when paired with a dTRE that activates them. The dotted red line corresponds to the distribution shown in Figure 2A. (G) Boxplots showing the percentage of promoters activated by dTREs of different regulatory natures.

**Supplemental Figure 3.**
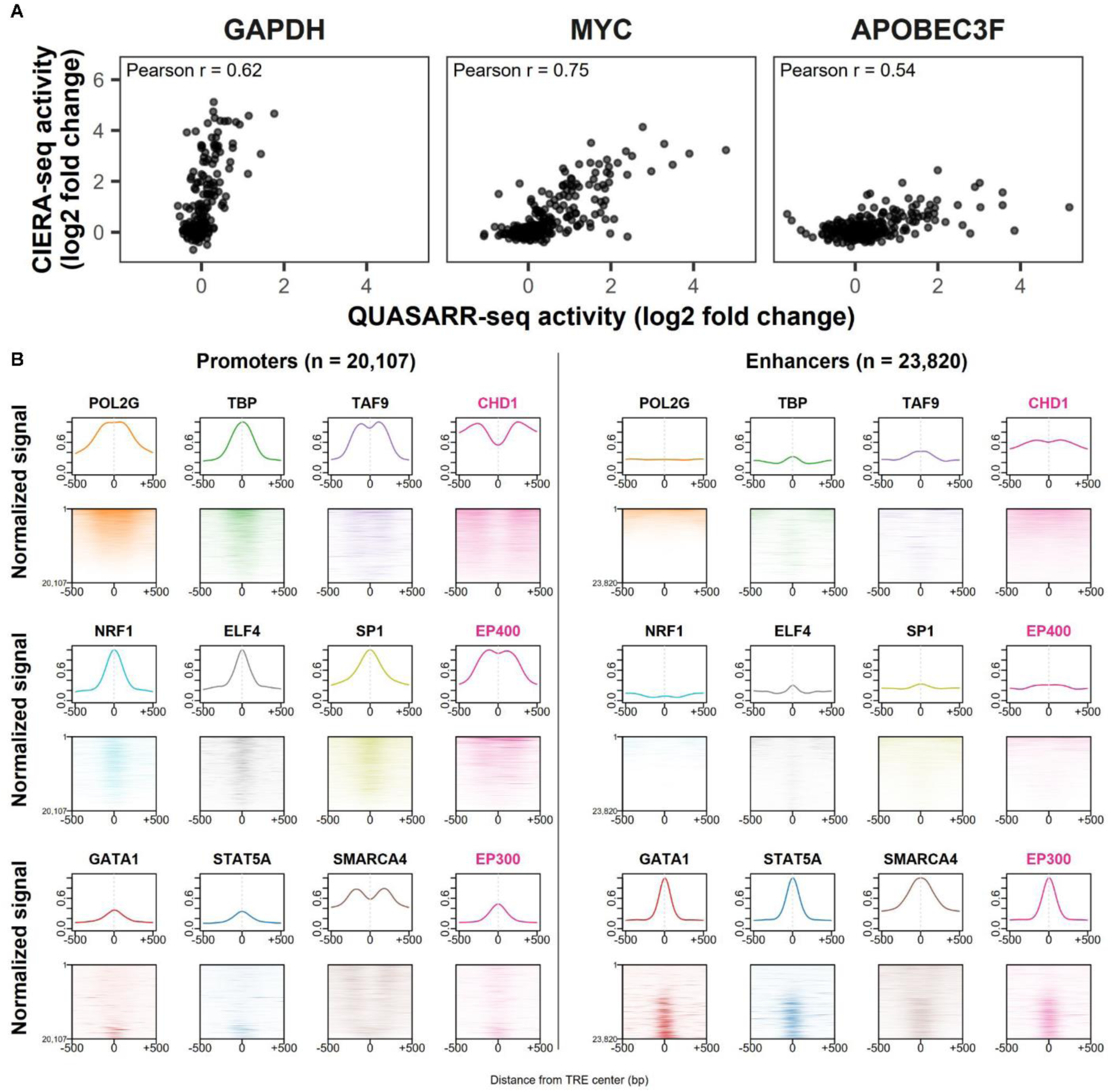
Comparison of CIERA-seq and QUASARR-seq and genome-wide transcription factor occupancy. (A) Correlation between plasmid-based enhancer activity measured by QUASARR-seq and CIERA-seq for the *GAPDH*, *MYC*, and *APOBEC3F* promoters. (B) Representative promoter-and enhancer-specific transcription factor and cofactor enrichment patterns derived f rom genome-wide TF ChIP-seq signals at promoters and enhancers.

**Supplemental Figure 4.**
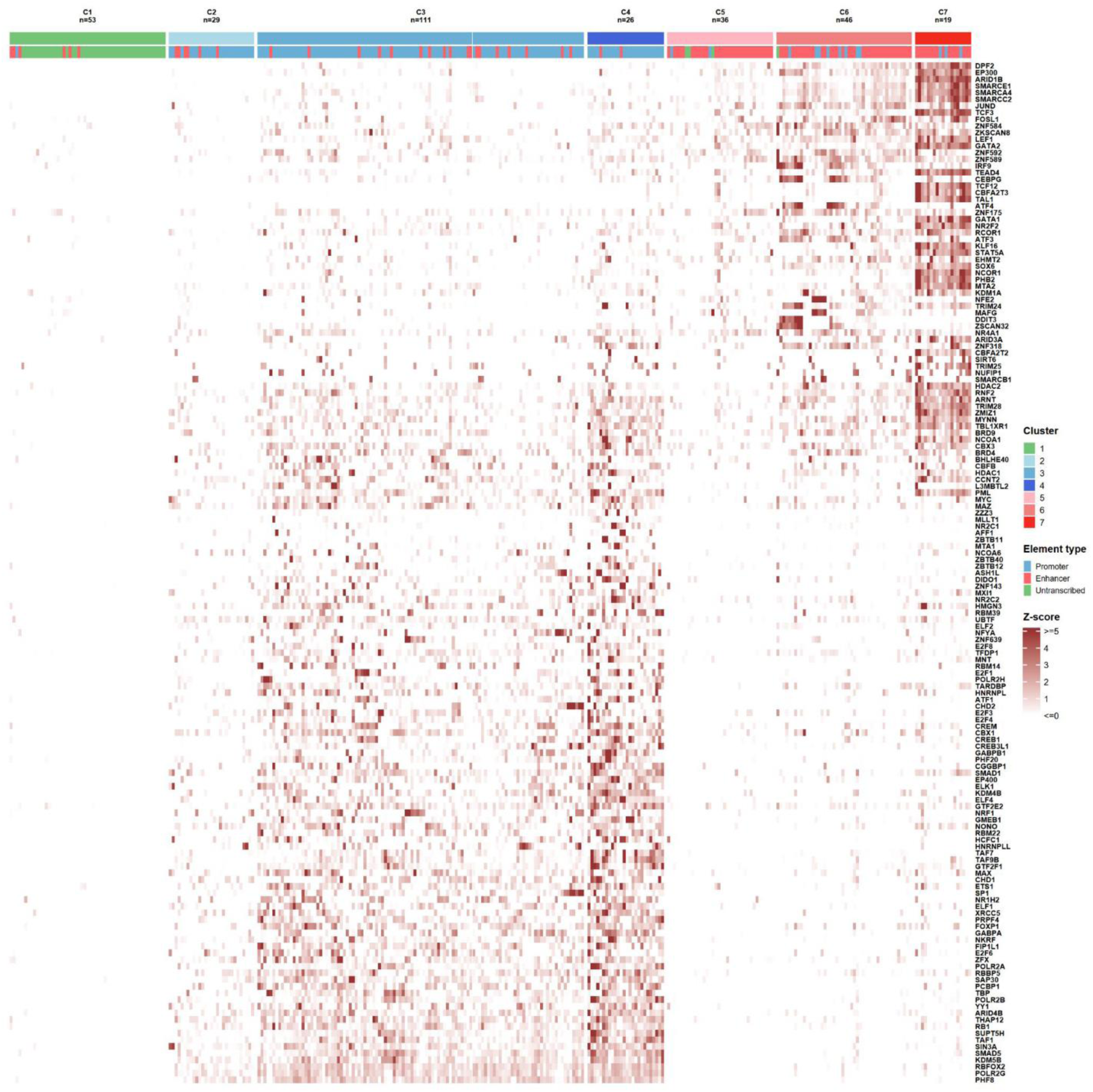
Transcription factor binding landscape across dTREs. Heatmap of ChIP–seq signals for all transcription factors that show differences in enrichment across dTREs. Columns represent individual dTREs, ordered by k-means clustering based on transcription factor binding profiles, and rows represent transcription factors.

**Supplemental Figure 5.**
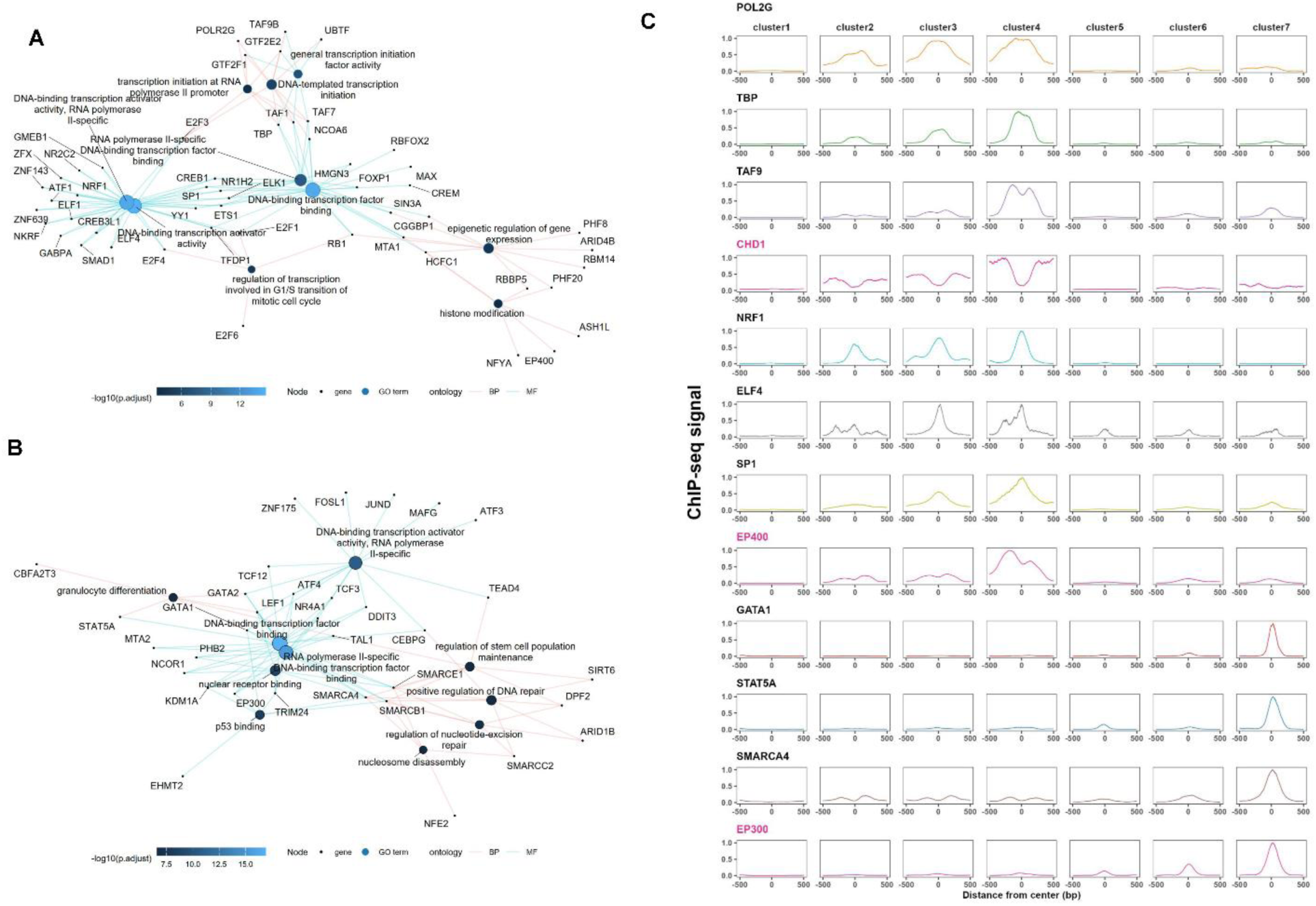
Gene Ontology enrichment and TF ChIP signals of promoter-and enhancer-biased transcription factors. (A) Network of gene ontology (GO) molecular function (MF) and biological process (BP) terms enriched among transcription fact ors showing higher enrichment at promoters than at enhancers. Solid small black dots represent transcription factors, and larger filled circles represent GO terms. Blue lines represent molecular function GO terms, and red lines represent biological process GO terms. (B) Network of gene ontology (GO) molecular function (MF) and biological process (BP) terms enriched among transcription fact ors showing higher enrichment at enhancers than at promoters. Solid black dots represent transcription factors, and larger filled circles represent GO terms. Blue lines represent molecular function GO terms, and red lines represent biological process GO terms. (C) Individual profiles of ChIP–seq signals for POL2G, TBP, TAF9, NRF1, ELF4, SP1, GATA1, STAT5A, CHD1, EP400, EP300, and SMARCA4 across transcriptional regulatory elements (dTREs) grouped by transcription factor–based k-means clusters.

**Supplemental Figure 6.**
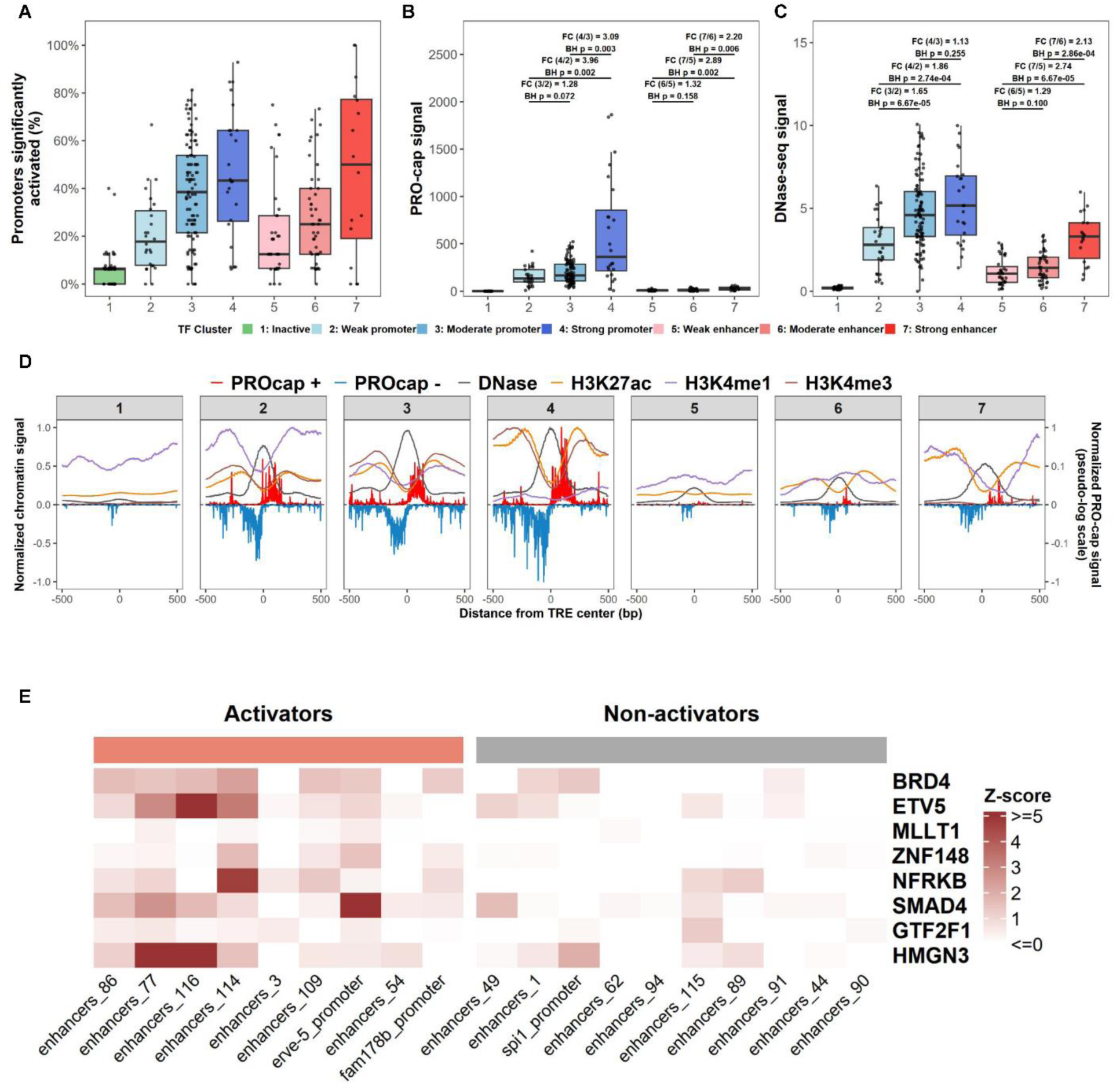
Activity and chromatin features of distal TRE clusters and TF enrichment in cluster 7. (A) Boxplots showing the percentage of promoters activated by dTREs f rom each transcription factor–based k-means cluster. (B) Boxplots of maximum PRO-cap signal across dTREs f rom each transcription factor–based k-means cluster; P values were calculated using the two-sided Welch tests adjusted using Benjamini–Hochberg procedure. (C) Boxplots of maximum DNase-seq signal across dTREs f rom each transcription factor–based k-means cluster; P values were calculated using the two-sided Welch tests adjusted using Benjamini–Hochberg procedure. (D) Profiles of PRO-cap, DNase-seq, and ChIP–seq signals for H3K27ac, H3K4me1, and H3K4me3 across dTREs f rom different transcription factor–based k-means clusters. ChIP-seq signals are plotted on the scale on the left side, and PRO-cap was plotted on a pseudo-log scale on the right side. (E) Heatmap of transcription factors differentially enriched across the cluster 7 dTREs that differ in their ability to activate the untranscribed target promoters.

**Supplemental Figure 7.**
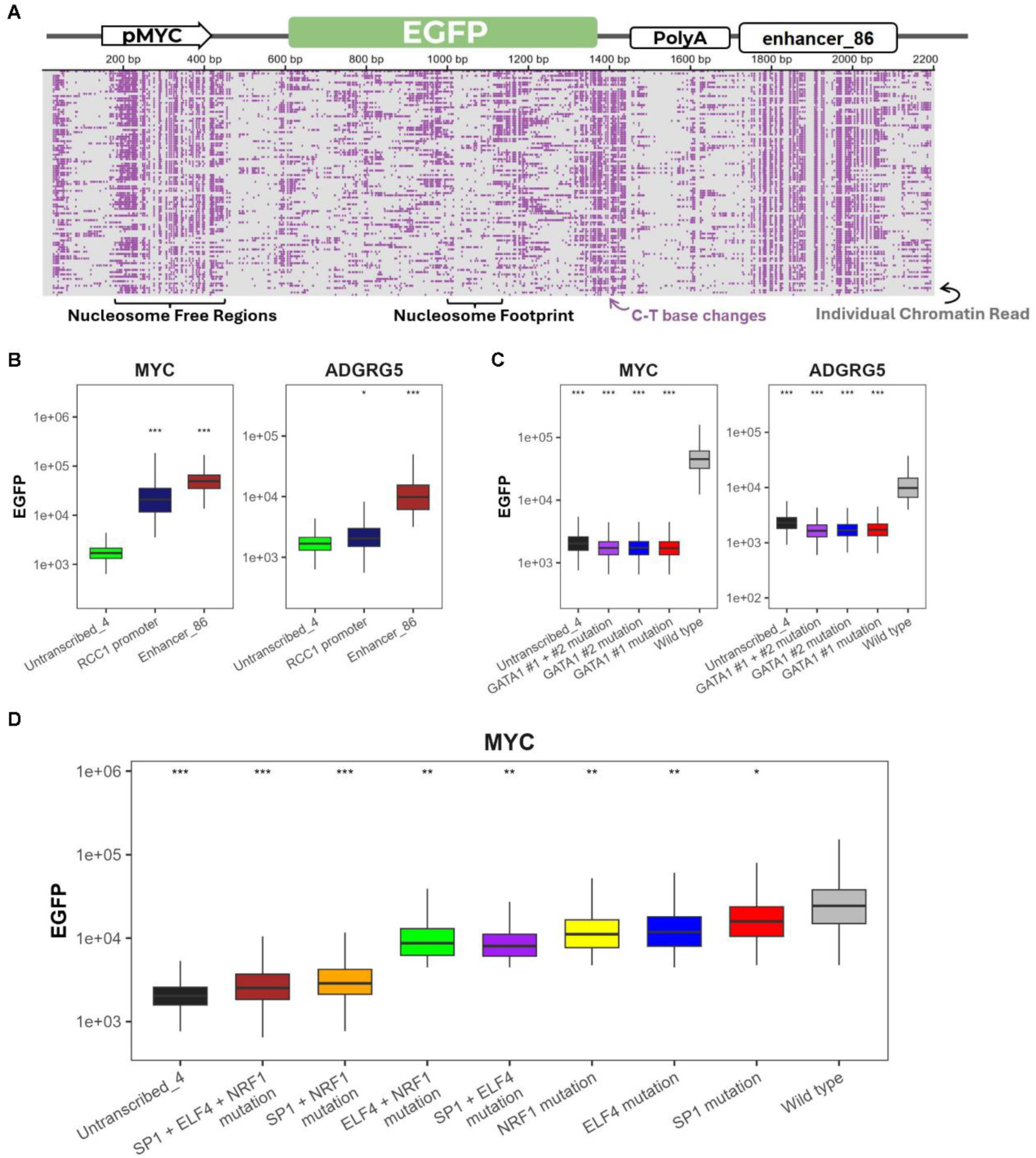
Deamination-based footprinting fibers and quantitative effects of dTRE class and motif mutations on promoter activity. (A) Single-fiber deamination patterns in cells containing the *MYC* promoter–enhancer_86 pair integrated at the landing-pad locus. (B) Boxplots of EGFP expression for cells harboring the *MYC* or *ADGRG5* target promoter paired with representative dTREs f rom different transcription factor–based clusters: untranscribed_4 dTRE (cluster 1), RCC1 promoter (cluster 4), and enhancer_86 (cluster 7). EGFP intensity is shown on the y-axis. Positive effects relative to the untranscribed control were assessed using Bonferroni-adjusted one-sided Welch lower confidence bounds. Effect thresholds of 0.1, 0.2, and 0.5 log10 units were met when the lower confidence bound exceeded the respective threshold. (C) Boxplots of EGFP expression for cells harboring the *MYC* or *ADGRG5* target promoter paired with the strong enhancer_86 (cluster 7) carrying different combinations of mutations in the two GATA1 motifs. Negative effects relative to the corresponding wild type were assessed using Bonferroni-adjusted one-sided Welch upper confidence bounds. Decreases exceeding 0.1, 0.2, and 0.5 log10 units were supported when the upper confidence bound was below −0.1, −0.2, and −0.5 log10 units, respectively. (D) Boxplots of EGFP expression for cells harboring the *MYC* target promoter paired with the RCC1 promoter (cluster 4) carrying different combinations of mutations in the SP1, NRF1, and ELF4 motifs. Negative effects relative to the corresponding wild type were assessed using Bonferroni-adjusted one-sided Welch upper confidence bounds. Decreases exceeding 0.1, 0.2, and 0.5 log10 units were supported when the upper confidence bound was below −0.1, −0.2, and −0.5 log10 units, respectively.

